# CHLOROPLAST UNUSUAL POSITIONING 1 is a new type of actin nucleation factor in plants

**DOI:** 10.1101/2020.01.14.905984

**Authors:** Sam-Geun Kong, Atsushi Shimada, Saku T. Kijima, Keiko Hirose, Kaoru Katoh, Jeongsu Ahn, Takeshi Higa, Akira Takano, Yuki Nakamura, Noriyuki Suetsugu, Daisuke Kohda, Taro Q. P. Uyeda, Masamitsu Wada

**Affiliations:** Department of Biology, Faculty of Sciences, Kyushu University, Fukuoka 812-8581, Japan; Department of Biological Sciences, College of Natural Sciences, Kongju National University, Chungnam 32588, Korea; Division of Structural Biology, Medical Institute of Bioregulation, Kyushu University, Fukuoka 812-8582, Japan; Biomedical Research Institute, National Institute of Advanced Industrial Science and Technology, Ibaraki 305-8562, Japan; Institute for Protein Research, Osaka University, Osaka 565-0871, Japan

**Keywords:** Actin, actin nucleator, Arabidopsis, blue light, chloroplast movement, CHUP1, formin, phototropin, profilin

## Abstract

Plants have evolved unique responses to fluctuating light conditions in their environment. One such response, chloroplast photorelocation movement, optimizes photosynthesis under weak light and prevents photodamage under strong light. CHLOROPLAST UNUSUAL POSITIONING 1 (CHUP1) plays a pivotal role in the light-responsive chloroplast movements, which are driven by dynamic reorganization of chloroplast actin (cp-actin) filaments. In this study, we demonstrated that fluorescently tagged CHUP1 colocalized and was coordinately reorganized with cp-actin filaments during chloroplast movement in *Arabidopsis thaliana*. The resulting asymmetric distribution of CHUP1 was reversibly regulated by the blue light receptor phototropin. X-ray crystallography indicated that the CHUP1 C-terminal domain shares structural similarity with the formin homology 2 (FH2) domain, although there is no sequence similarity between the two domains. The CHUP1 C-terminal domain stimulated actin polymerization in the presence of profilin. We conclude that CHUP1 is a novel, plant-specific actin nucleator that functions in cp-actin-based chloroplast movement.

**Highlights:**

1. Blue light changes the distribution pattern of CHUP1
2. Formin FH2 and CHUP1 C-terminal domains are structurally similar but not homologous
3. CHUP1 nucleates and severs actin filaments in vitro
4. CHUP1 is a novel, plant-specific actin nucleator for chloroplast movement

## INTRODUCTION

Plants, as sessile organisms, have evolved a variety of systems that allow them to adapt to fluctuations in their natural environments, such as changes in light intensity. Light is essential for photosynthesis but harmful when too strong. The chloroplast is a specialized organelle in plants and algae that converts solar energy into chemical energy through photosynthesis. In fluctuating light conditions, chloroplasts dynamically change their positions within the cell according to the quality, intensity, and direction of the incident light (Wada et al., 2003). Chloroplasts move toward weak light to capture as much energy as possible for photosynthesis (the accumulation response) (Gotoh et al., 2018; Zurzycki, 1955) but they rapidly move away from strong light to reduce light absorption (the avoidance response) (Kasahara et al., 2002; Zurzycki, 1957). Chloroplasts can move in any direction without rotating or rolling (Tsuboi and Wada, 2011; Tsuboi et al., 2009). When a small amount of light is shone on part of a cell, the cell’s chloroplasts move to the irradiated area (Kagawa and Wada, 1994). By contrast, when part of a chloroplast is irradiated directly with strong light, the chloroplast escapes from the light (Tsuboi and Wada, 2011).

Light-induced chloroplast movements are mediated by blue light receptors, specifically phototropins (phot1 and phot2 in seed plants) (Jarillo et al., 2001; Kagawa et al., 2001; Sakai et al., 2001). In *Arabidopsis thaliana*, phot1 functions only in the accumulation response (Sakai et al., 2001), whereas phot2 functions in both the accumulation and avoidance responses (Jarillo et al., 2001; Kagawa et al., 2001; Sakai et al., 2001). *phot2* mutants that are defective in the avoidance response are more susceptible than wild-type (WT) and *phot1* mutant plants to strong light-induced photodamage (Cazzaniga et al., 2013; Kasahara et al., 2002; Sztatelman et al., 2010). Thus, the avoidance response is essential for plant survival, especially under fluctuating light conditions (Wada et al., 2003). To control these movements, chloroplasts continuously monitor signals released from the photoreceptors (Tsuboi and Wada, 2013).

Chloroplast movement is mediated by dynamic reorganization of short actin filaments, called chloroplast actin (cp-actin) filaments, that are located between each chloroplast and the plasma membrane (Kadota et al., 2009). In response to strong blue light, rapid severing causes the cp-actin filaments to disappear transiently and then reappear on the leading edge of moving chloroplasts (Kadota et al., 2009; Kong et al., 2013a). phot2 induces the rapid severing that causes cp-actin depolymerization (Kong et al., 2013a). The distribution of cp-actin filaments around the chloroplast determines the speed and direction of the chloroplast’s movement; the larger the asymmetry in filament distribution, the faster the chloroplast moves (Kadota et al., 2009; Kong et al., 2013a)(reviews; Kong and Wada, 2011; Suetsugu and Wada, 2016).

CHLOROPLAST UNUSUAL POSITIONING 1 (CHUP1) functions in chloroplast positioning and movement, and *chup1* mutants are defective in chloroplast movement (Kasahara et al., 2002; Oikawa et al., 2003). CHUP1 has multiple functional domains, including an N-terminal hydrophobic region, a coiled-coil region, an actin-binding domain, a proline-rich region, and an uncharacterized C-terminal domain (Oikawa et al., 2003; Oikawa et al., 2008). CHUP1 is attached to the chloroplast outer envelope by its N-terminal hydrophobic domain (Oikawa et al., 2003; Oikawa et al., 2008). The actin binding domain of CHUP1 binds F-actin, and the C-terminal region binds profilin (Oikawa et al., 2003; Schmidt von Braun and Schleiff, 2008). Consistent with these activities, cp-actin filaments are not detected in *chup1* mutant cells, even though cytoplasmic actin filaments are clearly observed (Kadota et al., 2009; Kong et al., 2013a). The domain structure of CHUP1 and the *chup1* mutant phenotype suggest that CHUP1 functions in the regulation of cp-actin filaments. CHUP1 is highly conserved across a broad range of plant species. The protein is found in all plant lineages above the green algae (Charophytes) that show chloroplast photorelocation movement (Suetsugu and Wada, 2016), indicating that CHUP1 is the main player necessary for chloroplast movement in all plant species. In this study, we characterize the cellular and biochemical roles of CHUP1 and identify it as a novel plant-specific class of cp-actin nucleator functioning in chloroplast photorelocation movement.

## RESULTS

### CHUP1-YFP fusion protein is fully functional in transgenic plants

To examine CHUP1 function in the regulation of cp-actin filaments, we transformed a CHUP1-deficient Arabidopsis mutant (*chup1-3*) with CHUP1-fluorescent fusion proteins (CHUP1-YFP and CHUP1-tdTomato) driven by the native *CHUP1* promoter (Figure S1A). CHUP1-YFP was expressed at levels comparable to that of endogenous CHUP1 (Figure S1B). CHUP1-YFP expression successfully complemented the chloroplast positioning and photorelocation movement defects of the *chup1-3* mutant (Figures S1C, S1D). Under weak light (3.2 µmol m^−2^ s^−1^), chloroplasts accumulated on the periclinal wall in WT and CHUP1-YFP transgenic plants (C1Y) but not in *chup1-3* mutant plants. Under strong light conditions (25 and 60 µmol m^−2^ s^−1^), the chloroplast avoidance response was observed in both WT and C1Y transgenic plants but not in *chup1-3* mutant plants. The chloroplasts of *chup1-3* mutant plants were observed at the bottom of the cells under all light conditions, as previously described (Oikawa et al., 2003). Together, these data show that CHUP1-YFP in the transgenic plants is as fully functional as endogenous CHUP1 in WT plants.

### CHUP1 is reorganized on the chloroplast outer membrane in a blue light-dependent manner

Subcellular localization of CHUP1 under different light conditions was examined by confocal microscopy of palisade cells in rosette leaves of C1Y plants. When weak white light (20 µmol m^−2^ s^−1^) was applied for 2 h, CHUP1-YFP fluorescence was observed as small dots along the periphery of the plasma membrane side of the chloroplasts (Figures 1A middle panel, 1E). After dark adaptation for 1 h, the dots had aggregated to form longer and larger areas of fluorescence, mainly along the chloroplast periphery (Figure 1A left panel). When a part of a cell in a dark-adapted leaf was illuminated with strong blue light (458-nm laser, 5% output power) for several minutes, the dots of CHUP1-YFP fluorescence disappeared and were replaced by fluorescence distributed uniformly over the entire chloroplast envelope, including on the vacuole side (Figures 1D, 1E).

**Figure 1.**
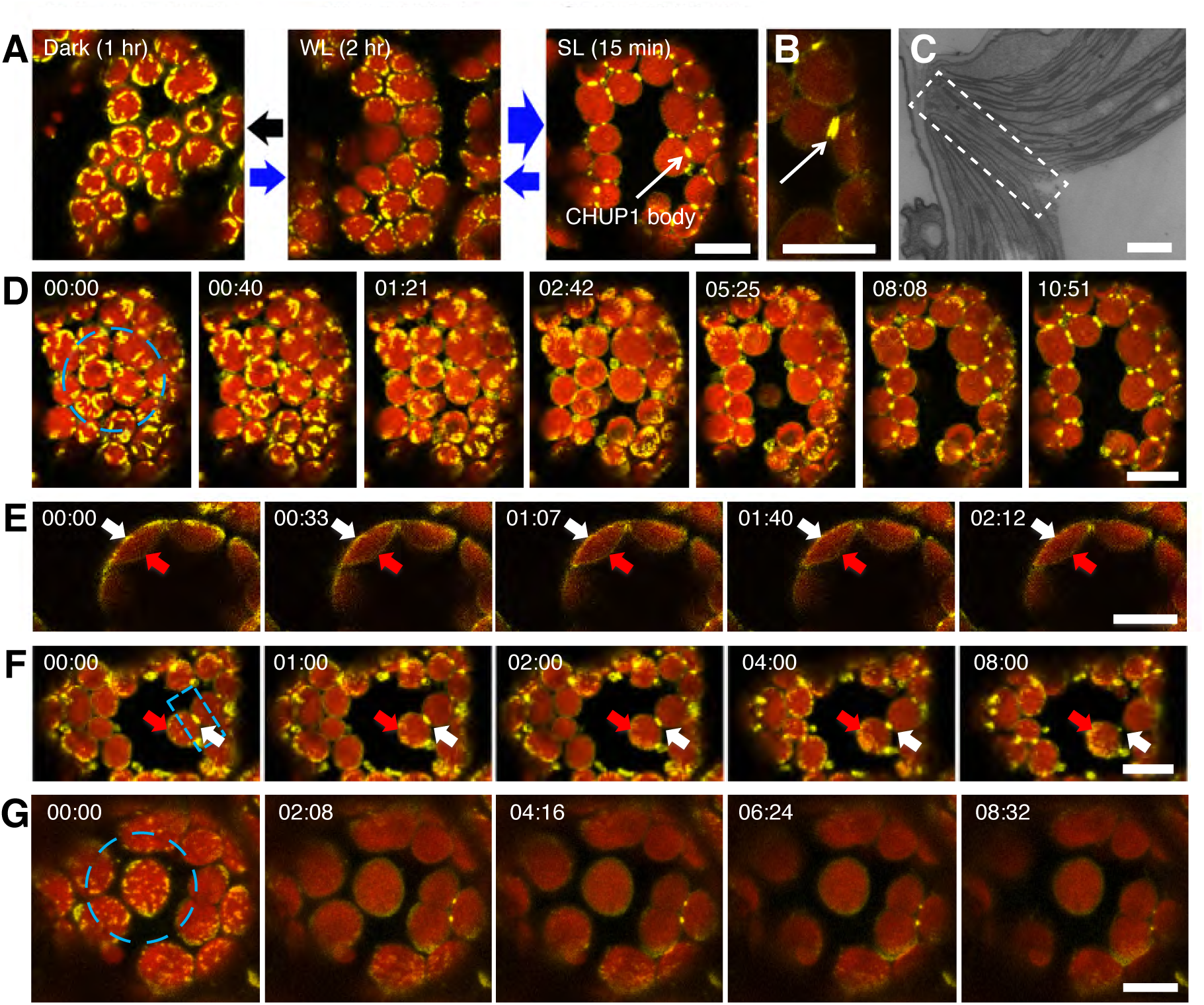
Reorganization of CHUP1-YFP on the chloroplast envelope in response to blue light. (A) Reversible changes in the distribution pattern of CHUP1-YFP under the indicated light conditions. CHUP1-YFP in the palisade cells of C1Y transgenic Arabidopsis was observed following incubation in darkness for 1 h (black arrow), in weak white light (WL; 20 µmol m^−2^ s^−1^) for 2 h (thin blue arrow), and in strong light (SL; 458-nm laser scan) for 15 min (thick blue arrow). (B) A CHUP1 body appears as two lines along the interface between two chloroplasts after 5 min of strong blue light irradiation. (C) TEM image of the region of contact (at the center of the rectangle) between two chloroplasts. (D) Blue light-dependent reorganization of CHUP1-YFP during a strong light-induced avoidance response followed by dark adaptation for 20 min. The dashed blue circle indicates the area that was irradiated. A time-lapse movie of this response can be seen in Movie S1. (E) Redistribution of CHUP1-YFP from the interface between the plasma membrane and chloroplast envelope (white arrows) to the entire chloroplast envelope, including the vacuole side (red arrows). The whole cell was irradiated continuously with strong blue light. (F) Redistribution of a CHUP1 body (white arrows) to small dots (red arrows) at the leading edge of a moving chloroplast. (G) Redistribution of CHUP1-YFP on an isolated irradiated chloroplast. The dotted CHUP1-YFP signal rapidly disappeared but no CHUP1 body was formed. A time-lapse movie of this response can be seen in Movie S2. The area irradiated with strong blue light is indicated as a dashed blue circle in D and G, and a dashed blue rectangle (10 µm x 5 µm) in F superimposed on the first image (00:00) of each series. Strong light was provided by 458-nm laser scans (5% output power). Time-lapse images were collected at 40-s, 33-s, 60-s, and 30-s intervals in D, E, F, and G, respectively, and are shown with false-color indicating YFP (yellow) and chlorophyll (red) fluorescence. The time (min:s) of image acquisition is shown in the upper left corner of each image. Scale bar = 500 nm in C, and 10 µm in all other panels.

The avoidance response of CHUP1-YFP plants was examined by illuminating a region of interest with a microbeam of strong light. After 15 min, chloroplast avoidance from the microbeam was clearly observed, and the dispersed CHUP1-YFP had accumulated at contact sites between chloroplasts, forming a specific structure, the ‘CHUP1 body’ (Figures 1A right panel, 1D; Movies S1). Sometimes a CHUP1 body appeared as two discrete lines of the same length, each belonging to one of the two adjacent chloroplasts (Figure 1B). Presumably, CHUP1 on the outer membrane of each chloroplast accumulated along the contact area and formed a CHUP1 body. We attempted to observe CHUP1 bodies in WT plants by electron microscopy. However, no notable structures were found where there was close contact between two chloroplasts’ outer membranes (Figure 1C). If a CHUP1 body was illuminated again with strong blue light, CHUP1-YFP fluorescence gradually decreased and small dots of fluorescence appeared at the leading edge of the moving chloroplast, suggesting that CHUP1 became redistributed on the chloroplast envelope (Figure 1F). If a chloroplast did not come into contact with other chloroplasts, no CHUP1 body was observed, and the chloroplast did not show directional movement (Figure 1G; Movie S2).

### Reorganization of CHUP1 is reversible and regulated in a blue light- and phototropin-dependent manner

Blue light influences not only chloroplast movements but also the distribution of CHUP1 on the chloroplast surface. Therefore, we examined whether phototropins are involved in the blue light-mediated regulation of CHUP1 localization. The C1Y line was crossed into *phot1chup1*, *phot2chup1*, and *phot1phot2chup1* mutant backgrounds to produce CHUP1-YFP/*phot1chup1*, CHUP1-YFP/*phot2chup1*, and CHUP1-YFP/*phot1phot2chup1* lines, respectively. The expression levels of CHUP1-YFP were similar in all lines (Figure 2A). When the avoidance response was induced in WT and *phot1* single mutant plants by partial cell irradiation with strong blue light (10 x 20 µm^2^ area), the chloroplasts moved out of the strong light-irradiated area and showed an asymmetric distribution of CHUP1-YFP fluorescence: higher intensity on the leading end and lower intensity on the rear end of the moving chloroplast (Figure 2B; Movies S3, S4). However, a partial reorganization of CHUP1-YFP was observed in the *phot2* single mutant (C1Y/*p2-1*) (Figure 2B; Movie S5). The dots of CHUP1-YFP fluorescence became smaller but never disappeared, and no CHUP1 body was formed (Figure 2B; Movie S5). No such reorganization of CHUP1-YFP was observed in *phot1phot2* double mutant cells (C1Y/*p1-5p2-1*) (Figure 2B; Movie S6). By contrast, the dots of CHUP1-YFP fluorescence disappeared normally in the *phot1* single mutant (C1Y/*p1-5*), indicating that phot2 alone is sufficient for CHUP1 reorganization and that phot1 is partially functional in the regulation of CHUP1 reorganization. Therefore, CHUP1 localization was dynamically and reversibly regulated in a blue light- and phototropin-dependent manner.

**Figure 2.**
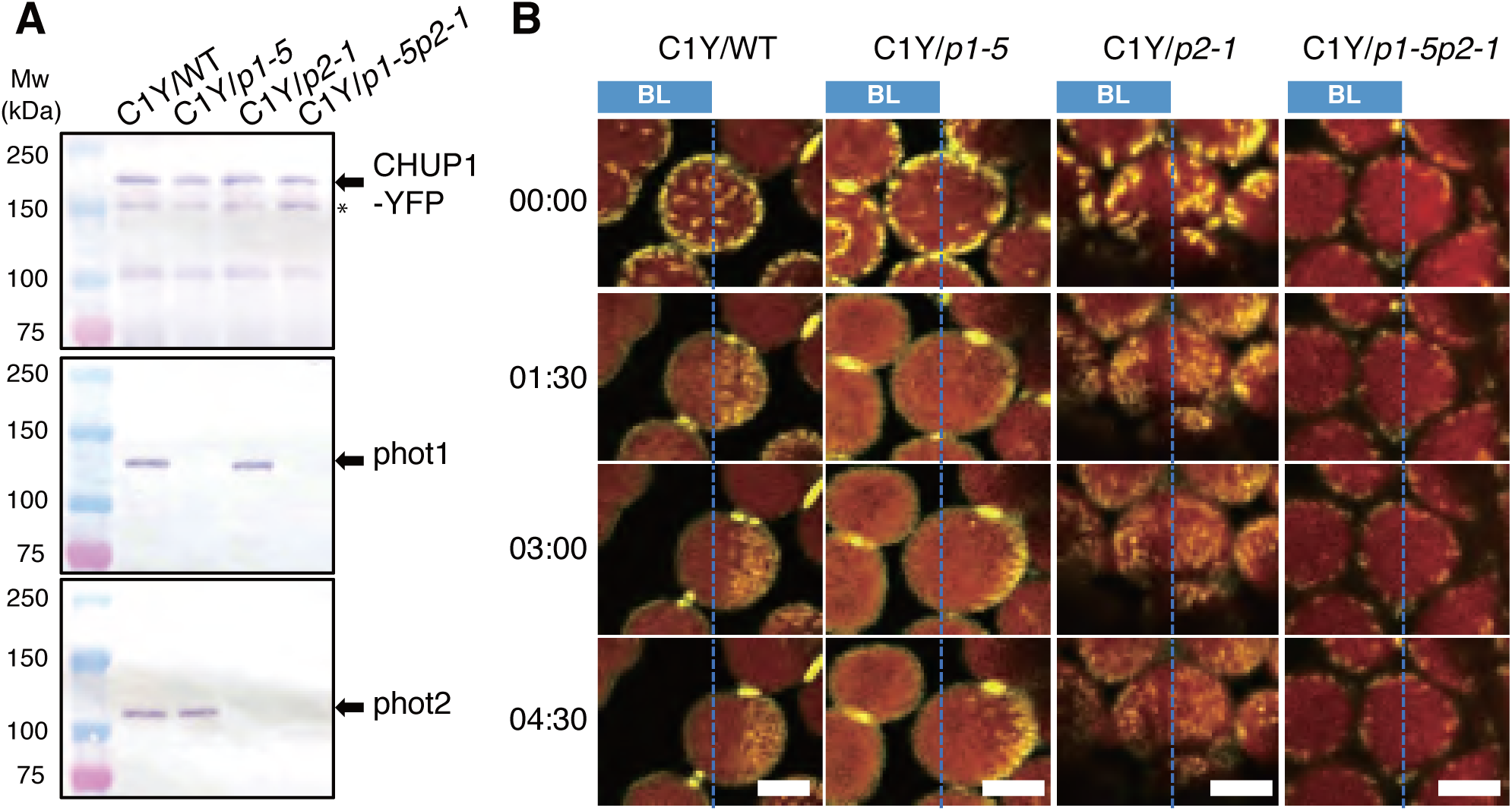
Identification of the main photoreceptor regulating CHUP1 localization. (A) Immunoblot analysis of CHUP1-YFP expressing transgenic plants in the WT, *phot1*, *phot2*, and *phot1phot2* mutant backgrounds (C1Y/WT, C1Y/*p1-5*, C1Y/*p2-1*, C1Y/*p1-5p2-1*). Protein samples (50 µg) were separated on 7.5% SDS-PAGE gels and probed with anti-CHUP1 (top panel), anti-PHOT1 (middle), and anti-PHOT2 (bottom) polyclonal antibodies. Arrows indicate CHUP1-Y, phot1, and phot2. * indicates truncated CHUP1-Y. (B) Series of images at the indicated time points (min:s) show reorganization of CHUP1-YFP in strong blue light in WT and *phot*-deficient mutant cells. In each image, the region to the left of the blue dotted line was irradiated with strong blue light by 458-nm laser scans at 30-s intervals to induce the avoidance response. Scale bars = 5 µm. Time-lapse movies of these responses can be seen in Movies S3–S6.

### CHUP1 colocalizes and is reorganized coordinately with cp-actin filaments

We visualized cp-actin filaments in Arabidopsis transgenic plants expressing GFP fused with the actin binding domain of mouse Talin (GFP-mTalin). In these plants, we observed an asymmetric distribution of cp-actin filaments in the region between the plasma membrane and the moving chloroplasts (Kadota et al., 2009; Kong et al., 2013a). The distribution pattern and dynamics of CHUP1 were similar to those of the cp-actin filaments (Figure 3E). Therefore, we studied the relationship between cp-actin filaments and CHUP1 during chloroplast movements in a transgenic line expressing CHUP1-tdTomato and GFP-mTalin (CHUP1-tdTomato x GFP-mTalin line). CHUP1 and cp-actin filaments were co-localized as expected (Figures 3A, 3B, 3F). Both CHUP1 and cp-actin filaments disappeared rapidly in response to strong blue light and reappeared at the front ends of moving chloroplasts (Figure 3F, Movie S7), as previously observed for cp-actin filaments (Kadota et al., 2009; Kong et al., 2013a).

**Figure 3.**
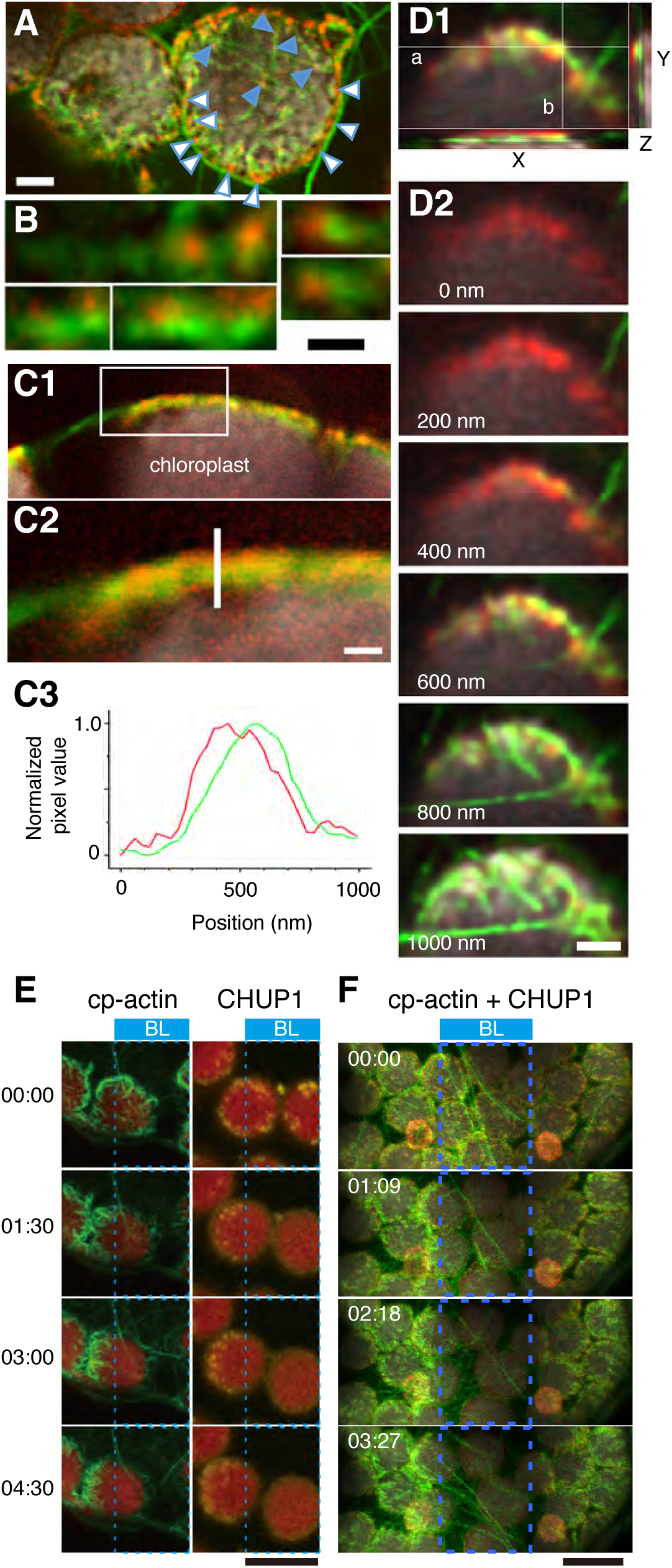
Association of CHUP1 with cp-actin filaments. (A) Top view of chloroplasts facing the plasma membrane of a periclinal wall of the palisade cells. Cp-actin filaments were visualized with GFP-mTalin (green) and CHUP1 with tdTomato (red) in a CHUP1-tdTomato × GFP-mTalin line. CHUP1 was mostly present on the chloroplast periphery. White arrowheads indicate CHUP1 attached to long cytoplasmic actin filaments. Blue arrowheads indicate CHUP1 localized at the tips or along the sides of cp-actin filaments. (B) Magnified views of CHUP1 localized close to cp-actin filaments. (C) Distribution of CHUP1 and cp-actin filaments within the region between the plasma membrane and the chloroplast envelope. CHUP1 localized to the plasma membrane side and cp-actin filaments along the chloroplast side. C1: an optical slice of a side view of a chloroplast next to the plasma membrane along an anticlinal wall. C2: an enlarged image of the area within the rectangle in C1. C3: fluorescence intensity profiles of CHUP1-tdTomato (red) and cp-actin filaments (green) along the white line in C2 from the plasma membrane side (0, top of the line) to the inside (1000) of the chloroplast. (D) Separate distributions of CHUP1 at the plasma membrane side and cp-actin filaments on the chloroplast side. Top views of part of a chloroplast are shown. D1: A Z-series of six confocal images of fluorescence of CHUP1-tdTomato (red) and GFP-mTalin (green) taken with 200-nm steps from the plasma membrane side to the chloroplast side and superimposed on an X-Y plane using the maximum intensity projection method. In the X-Z plane at white line a and the Y-Z plane at white line b, the CHUP1-tdTomato and GFP-mTalin fluorescence for the entire Z-series was compiled. This confirmed that CHUP1 is on the plasma membrane side and cp-actin filaments are on the chloroplast side. The individual images are shown in D2. Scale bars are 1 µm in A and D, and 500 nm in B and C. (E) Asymmetric distributions of cp-actin filaments (left panels) and CHUP1 (right panels) on moving chloroplasts. Time-lapse images were collected at 30-s intervals for 4 min 30 s under strong blue light irradiation using 458-nm laser scans (blue rectangles indicate region of interest [roi]) in the intervals between image acquisitions. Image acquisition time stamps (min:s) are shown. Scale bar = 10 µm. (F) Reorganization of CHUP1 and cp-actin filaments on moving chloroplasts in a CHUP1-tdTomato × GFP-mTalin line. Time-lapse images were collected at the indicated time points (min:s). The other details are the same as in (E), except that images were acquired at 34-s intervals for 3 min 27 s. Scale bar = 10 µm. A time-lapse movie of this response is shown in M**ovie S7**.

CHUP1-tdTomato fluorescence appeared as dots mainly around the periphery of the plasma membrane side of the chloroplasts (Figures 3A, 3F). In many cases, the CHUP1 dots appeared to be connected to one end or to the side of cp-actin filaments (Figures 3A, 3B, 3F). Optical sectioning demonstrated that CHUP1-tdTomato was observed close to the plasma membrane, but not to the chloroplasts (Figures 3C, 3D). This observation was confirmed by fluorescence intensity analysis of the interface between the plasma membrane and a chloroplast (Figure 3C). Long cytoplasmic actin filaments were frequently observed running along the chloroplast periphery and attaching to the CHUP1 dots there (Figure 3A).

### The CHUP1 C-terminal fragment has structural homology to formin homology-2 domains

To gain insight into CHUP1 function, we determined the crystal structure of the CHUP1 C-terminal fragment, CHUP1_756-982 (Table S1). The overall structure of CHUP1_756-982 has an elongated shape consisting entirely of α-helices. Intriguingly, the three-helix bundle at the center of the C-terminal domain is reminiscent of the formin homology-2 (FH2) domains of all well-characterized actin nucleators (Figures 4A, 4B; Shimada et al., 2004; Xu et al., 2004). Although there is no obvious amino acid sequence similarity between the two domains and the corresponding region of the FH2 sequence is non-contiguous, the two structures are superimposable with the root-mean-square deviation of 1.82 Å over the 52 C_α_ atoms.

**Figure 4.**
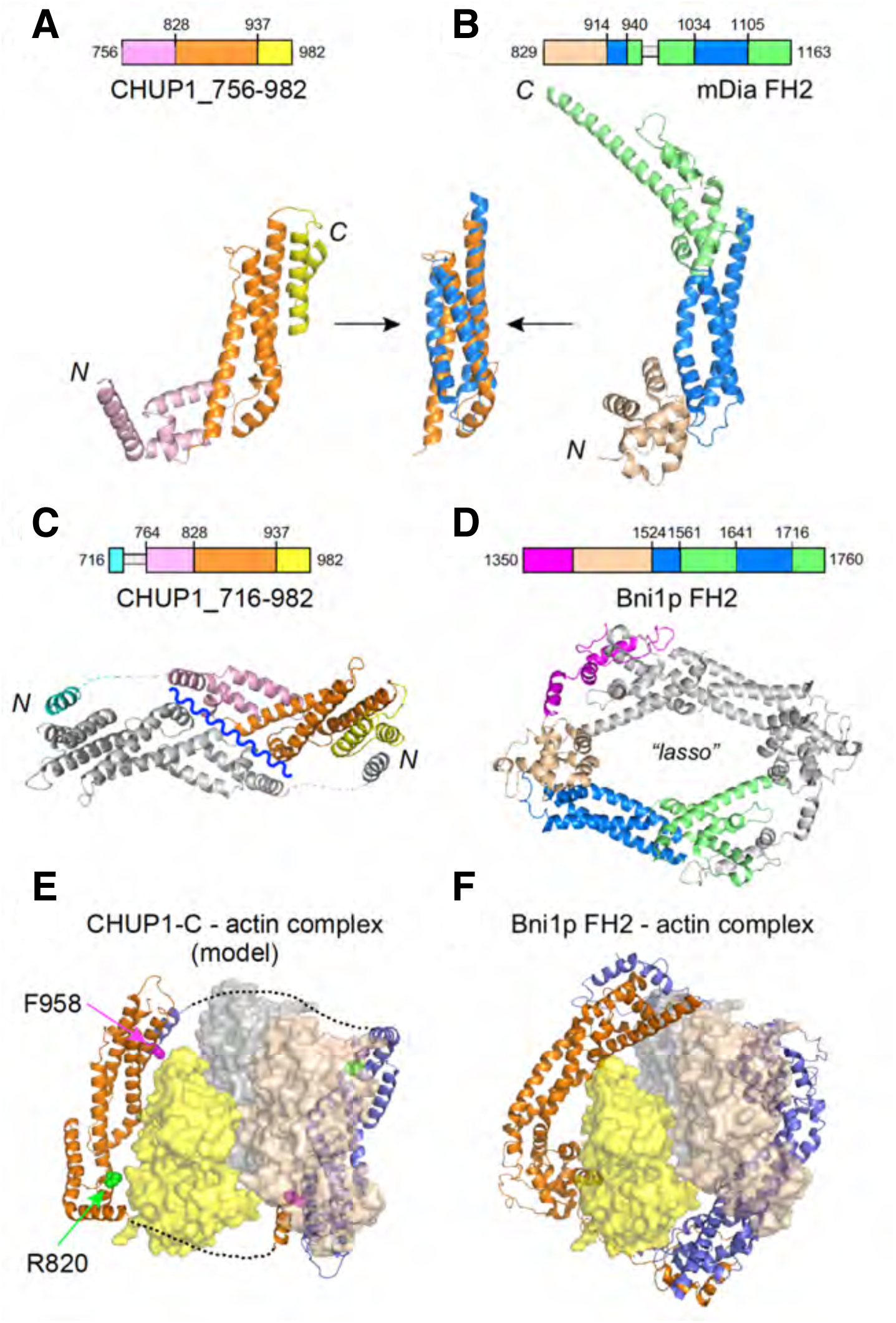
Three-dimensional structures of the C-terminal fragments of CHUP1. (A) Crystal structure of CHUP1_756-982. (B) Crystal structure of the mDia1 FH2 domain (PDB 1V9D). The central three-helix bundles of CHUP1_756-982 and FH2 are superimposed in the middle. (C) Crystal structure of CHUP1_716-982. This fragment adopts a closed dimeric structure, which might open along the wavy line to become a functional dimer. (D) Functional dimer form of Bni1p FH2 (PDB 1UX5). (E) Model of the CHUP1_716-982 dimer and actin complex. The positions of R820 (green) and F958 (magenta) are shown. (F) Crystal structure of the Bni1p FH2 dimer and actin complex (PDB 1Y64). Loops that are missing from the crystal structures because of poor electron density are indicated by dashed lines in the structures and by thin gray boxes in the amino acid sequences.

The functional form of the FH2 domain is a lasso-shaped dimer, in which the two subunits interact at both ends of their elongated structures (Figure 4D; Xu et al., 2004). The CHUP1 C-terminal fragment used in the crystallization described above did not contain the sequence corresponding to the N-terminal interacting site of the FH2 domain. Therefore, we extended the original CHUP1 sequence by 40 residues to the N-terminus and determined the crystal structure of CHUP1_716-982 (Table S1). The newly added N-terminal portion forms a short α-helix and connects with another CHUP1 molecule in the crystal (Figure 4C). The 30-residue segment that connects with the short, N-terminal α-helix and CHUP1_756-982 had no clear electron density (broken lines in Figure 4C). We speculate that this segment serves as a flexible tether, allowing a functional dimer to form (Figure 4E) in a manner similar to the FH2 domain–actin complex (Figure 4F).

### CHUP1-C interacts with the barbed ends and sides of actin filaments

To examine how CHUP1 interacts with actin filaments, a preliminary cosedimentation assay was performed using CHUP1_611-1004, hereafter called CHUP1-C, and rabbit skeletal muscle actin filaments (Figure S2A and Table S2). When 1 µM CHUP1-C was allowed to interact with 10 µM rabbit skeletal muscle actin filaments, a fraction of CHUP1-C co-sedimented with the actin filaments following ultracentrifugation. When the actin filaments were sheared by vigorous pipetting prior to the addition of CHUP1-C, the co-sedimented fraction of CHUP1-C increased from 4.7% to 10.4%, suggesting that CHUP1-C interacts with ends of actin filaments. By contrast, fragmentation of the actin filaments with gelsolin did not increase the amount of CHUP1-C that co-sedimented. Because gelsolin remains bound to the barbed ends of actin filaments after severing (Sun et al., 1999), this result suggested that CHUP1-C binds to barbed ends, but not to pointed ends, of actin filaments. Nonetheless, the fact that some CHUP1-C bound to gelsolin-treated actin filaments suggested that CHUP1-C can bind along the sides of actin filaments as well, which was confirmed by electron microscopy, as described in the next section. The above experiments were performed in the presence of Ca^2+^, which was required to activate gelsolin. However, interactions of CHUP1-C with actin filaments did not depend on Ca^2+^, since chelating Ca^2+^ with excess EGTA did not change the amount of co-sedimented CHUP1-C.

Fluorescence microscopy observation of actin filaments after staining with rhodamine-phalloidin (Figure S2B) confirmed that shearing by vigorous pipetting decreased the average filament length. Interestingly, actin filaments that had interacted with 1 µM CHUP1-C, without shearing or treatment with gelsolin, were also significantly shorter than the control. This result suggested that CHUP1-C has actin filament severing activity, which may be related to the binding of CHUP1-C along the sides of actin filaments.

### CHUP1 is an actin nucleator/severer

We next tested whether the CHUP1-C fragment, which includes the polyproline sequences and the FH2-like domain (Figure 6A), binds to the barbed ends of actin filaments and promotes polymerization of actin that is complexed with profilin, a hallmark activity of formin family proteins.

**Figure 5.**
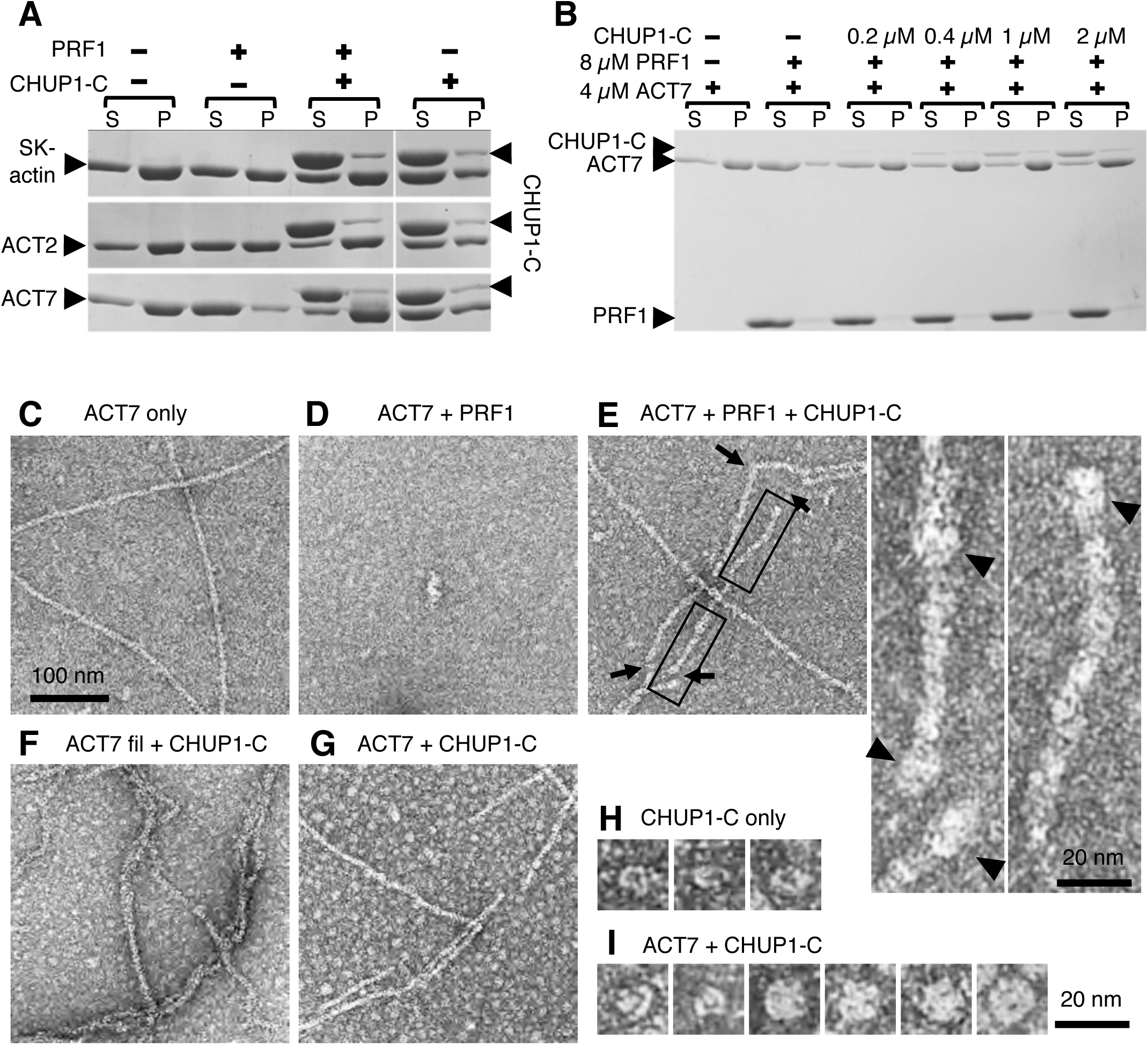
Interaction of CHUP1-C with actin filaments in vitro. (A) Ultracentrifugation assays of the effects of profilin and CHUP1-C on actin polymerization. Rabbit skeletal muscle actin or Arabidopsis ACT2 or ACT7 was allowed to polymerize in F-buffer in the presence or absence of 8 µM PRF1 and 4 µM CHUP1-C. After ultracentrifugation, the supernatant (S) and pellet (P) fractions were separately run on SDS-PAGE. (B) Similar to A, except that only ACT7 was used and the concentration of CHUP1-C was varied from 0 to 2 µM. (C–I) Electron micrographs of actin polymerized in the presence or absence of profilin and CHUP1-C. In (C–E), 4 µM ACT7 was allowed to polymerize in F-buffer in the absence of PRF1 or CHUP1-C (C), in the presence of 4 µM PRF1 (D), and in the presence of 4 µM PRF1 and 0.4 µM CHUP1-C (E). Enlarged views of the boxed regions in (E) are shown on the right. Arrows show sharp bends and gaps along the filaments. (F) ACT7 (4 µM) was polymerized in F-buffer for 20 min and then incubated with 0.4 µM CHUP1-C for 1-2 min. (G) ACT7 (4 µM) was allowed to polymerize in F-buffer in the presence of 0.4 µM CHUP1-C for 20 min. (H) CHUP1-C (0.4 µM) was incubated in F-buffer for 5 min and then diluted to 0.2 µM. Particles observed in the sample prepared as in (G) are shown (I). All samples except for (H) were diluted to 0.7 µM ACT7 in EM buffer, and then negatively stained. The magnification in (D–G) is the same as in (C) (scale bar = 100 nm). The enlarged insets in (E) are the same magnification as the particles in (H, I) (scale bar = 20 nm).

**Figure 6.**
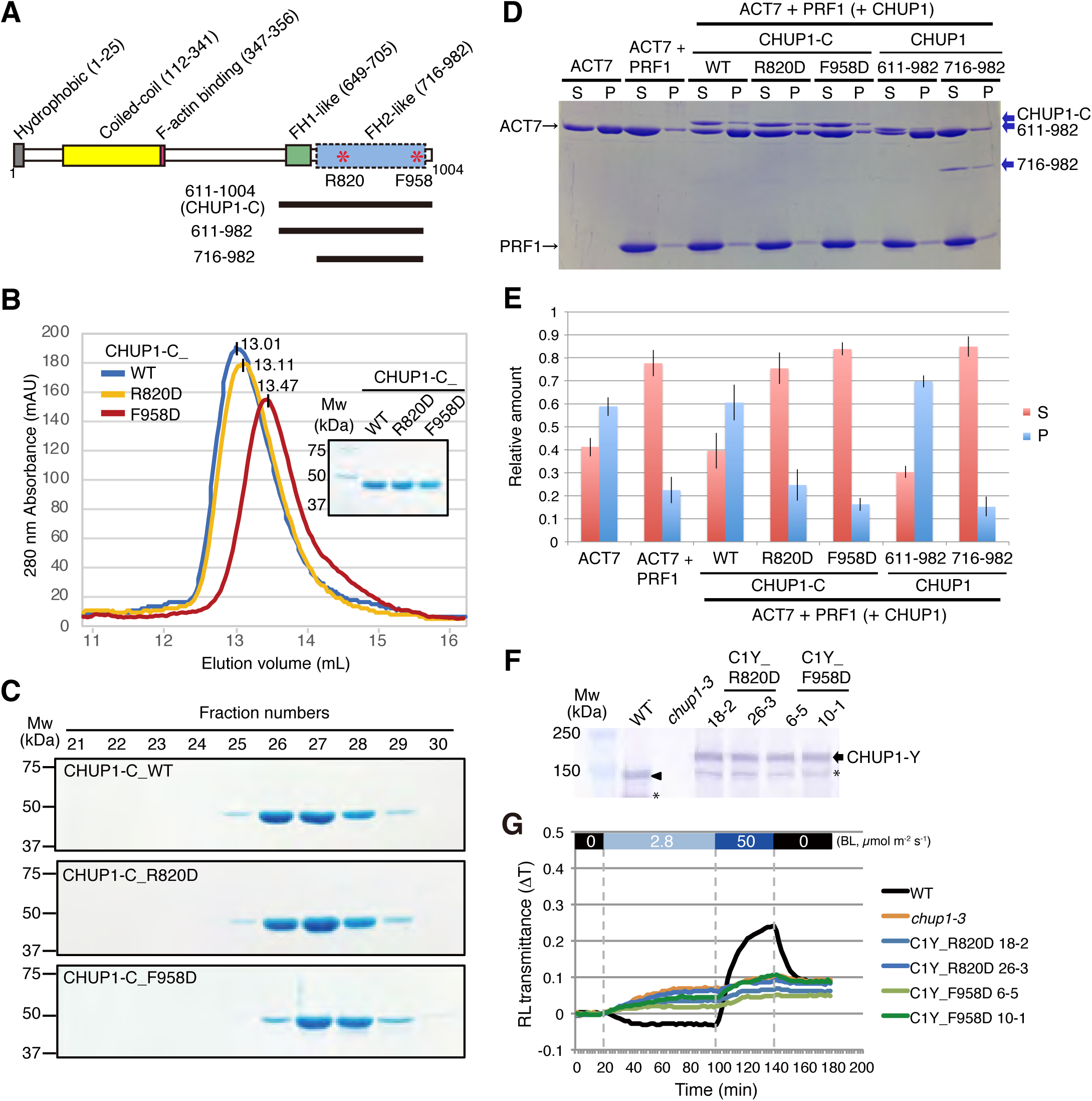
Effects of mutations at R820 and F958 of CHUP1-C on the activity of CHUP1. (A) A schematic illustration of CHUP1 protein showing the functional domains: the hydrophobic region, the coiled-coil domain, the F-actin binding domain, the proline-rich FH1-like domain, and the FH2-like domain. Asterisks indicate R820, which is involved in the association with actin, and F958, which is involved in the dimerization of the FH2-like domain. The positions of the 611–1004 fragment (CHUP1-C) and the 611–982 and 716–982 deletion mutations are indicated by lines below the structure. (B–C) The effect of point mutations R820D and F958D on CHUP1-C dimer structure. (B) The elution profiles of CHUP1-C_WT, R820D, and F958D recombinant proteins were obtained by gel filtration chromatography using a Superdex 200 10/300 column. (C) The protein profiles were confirmed using 10% SDS-PAGE gels. (D–E) Ultracentrifugation assay of actin polymerization. ACT7 (4 µM) was allowed to polymerize in F-buffer containing 8 µM PRF1 and 0.4 µM WT or mutant CHUP1-C. (D) A representative SDS-PAGE gel of supernatant (S) and pellet (P) fractions after ultracentrifugation. (E) A bar graph showing the fraction of ACT7 in the pellet and the supernatant under each condition, with error bars showing SD (N = 3). (F) Immunoblot analysis showing the expression levels of CHUP1-YFP_R820D and CHUP1-YFP_F985D in C1Y_R820D (18-2 and 26-3) and C1Y_F958D (6-5 and 10-1) transgenic plants. The arrowhead and arrow indicate CHUP1 and CHUP1-Y, respectively. * indicates truncated CHUP1 and CHUP1-Y. The details are the same as in Figure 2A. (G) Chloroplast photorelocation movement in WT, *chup1* mutant (*chup1-3*), and C1Y_R820D and C1Y_F985D transgenic plants in a *chup1* background. The red light (RL) transmittance in rosette leaves was monitored under different intensities of blue light (BL; 0, 2.8, and 50 µmol m^−2^ s^−1^) for the indicated periods.

To assess the polymerization promoting activities of CHUP1-C, we used ACT2 and ACT7, two major isoforms of Arabidopsis vegetative actin, in addition to skeletal muscle actin. As we reported previously (Kijima et al., 2016), ultracentrifugation assays demonstrated that prior addition of excess Arabidopsis profilin (PRF1) strongly inhibited salt-induced spontaneous polymerization of ACT7, while polymerization of ACT2 or skeletal muscle actin was only partially inhibited by PRF1 (Figure 5A). This inhibition was fully reversed by the addition of 4 µM CHUP1-C in the case of ACT2 and ACT7, but only partially reversed in the case of skeletal muscle actin. When CHUP1-C was added to G-actin in the absence of PRF1, polymerization was partially inhibited (Figures 5A, S2C). These results suggest that CHUP1-C has a potent formin-like ability to polymerize autologous actin complexed with autologous profilin, but has weaker activity with heterologous muscle actin (Figure 5A).

Next, we further examined the ability of CHUP1-C to promote polymerization of actin complexed with PRF1 using ACT7. Polymerization promoting activity was evident when 0.2 µM CHUP1-C was added to 4 µM ACT7 and 8 µM PRF1, and the activity was saturated at 0.4 µM (Figure 5B). The ability of CHUP1-C to promote polymerization of ACT7 in the presence of PRF1 was confirmed by kinetic experiments (Figure S4) and by electron microscopy observation after negative staining with uranyl acetate (Figures 5D, 5E). These results are consistent with the idea that CHUP1-C has a formin-like actin-nucleating activity.

We noted that unlike control ACT7 filaments (Figure 5C), ACT7 filaments polymerized in the presence of PRF1 and CHUP1-C had frequent sharp bends, and some filaments showed gaps along otherwise continuous filaments (arrows in Figure 5E). These bends and gaps were probably caused by weakened interactions between actin molecules along the filaments, which were disrupted upon landing on the carbon film during sample preparation. Similarly, we observed contorted filaments with frequent gaps when CHUP1-C was added to filaments of ACT7 (Figures 5F, S5) or when ACT7 was allowed to polymerize in the presence of CHUP1-C but in the absence of PRF1 (Figure 5G). Similar electron microscopy images were obtained with muscle actin filaments in the presence of CHUP1-C (Figure S3).

At higher magnifications, blob-like structures were sometimes observed at the ends and along the sides of ACT7 filaments in the presence of CHUP1-C (arrowheads in Figure 5E). These blobs resembled the particles we observed in the background of ACT7-CHUP1-C mixtures and are probably CHUP1-C molecules or oligomers associated with actin (Figure 5I). These results support the hypothesis that CHUP1-C binds not only to the ends of actin filaments but also along their sides to sever them.

### Experimental support for a hypothetical model of the CHUP1_716-982-actin complex

Based on our results, we constructed a hypothetical model of the CHUP1_716-982-actin complex with the assumption that CHUP1_716-982 forms a dimer that is structurally and functionally similar to FH2 (Figure 4E). To test the validity of this model, we selected two amino acid positions for mutagenesis. In the hypothetical complex, the R820 residue is positioned close to the contact sites with the actin molecules, and substitution at this position would be expected to reduce actin binding. F958 is located near the C-terminal end of the CHUP1_716-982 fragment and interacts with the short α-helix in the N-terminally extended region. An amino acid substitution at F958 is thus expected to disrupt formation of a functional dimer.

We used size-exclusion chromatography to measure the ability of mutagenized CHUP1-C domains to form complexes. CHUP1-C with a F958D point mutation (CHUP1-C_F958D) eluted more slowly than CHUP1-WT and CHUP1-C with a R820D point mutation (CHUP1-C_R820D), consistent with our prediction that F958 is involved in multimer formation (Figures 6B, 6C). If dimerization is essential for CHUP1-C activity, as is the case for formins, the CHUP1-C_F958D mutant would be expected to be unable to promote polymerization of ACT7 in the presence of PRF1. This prediction was confirmed by sedimentation experiments (Figures 6D, 6E). Likewise, CHUP1-C_R820D lacked the ability to promote polymerization of ACT7 in the presence of PRF1 (Figures 6D, 6E).

The physiological functions of R820 and F958 were further confirmed in transgenic plants expressing CHUP1-YFP with R820D or F958D substitutions in a *chup1-3* background (Figures 6F, 6G). When chloroplast movements in the two transgenic lines were examined by a red-light transmittance assay, neither the accumulation response nor the avoidance response was detected (Figure 6G). Thus, the chloroplast movement phenotype of the *chup1-3* mutant was not rescued in the two transgenic plants, and both R820 and F958 are essential for CHUP1 function.

The region between amino acid residues 611 and 716 includes polyproline sequences, which correspond to the formin homology-1 (FH1) domain that was implicated in profilin binding (Schmidt von Braun and Schleiff, 2008). Removal of this polyproline region, together with a C-terminal truncation at 982, completely inhibited the ability of CHUP1 to promote polymerization of ACT7 in the presence of PRF1 (Figures 6D, 6E). However, the C-terminal truncation at 982 on its own had no noticeable effect on the actin-polymerizing activity (Figures 6D, 6E). Thus, the polyproline sequence is also important for CHUP1-mediated polymerization of actin bound to profilin.

Based on the results of the structural, biochemical, and microscopy analyses described above, we conclude that CHUP1-C possesses formin-like activity that promotes actin polymerization in the presence of profilin and severs actin filaments.

## DISCUSSION

### CHUP1 is a newly identified plant-specific class of actin nucleator

Our study shows that CHUP1 is a formin-like actin nucleator that polymerizes cp-actin (Figure 5). Although the crystal structure of the CHUP1 C-terminal fragment, CHUP1_756-982, has some structural homology to the FH2 domain of formin (Figure 4), there is no detectable similarity between the two amino acid sequences. From the structural model of the CHUP1_716-982-actin complex, we predicted that F958 could be important for dimerization of CHUP1 and that R820 could be part of a potential actin-binding site (Figure 4E). These residues were confirmed to be essential for actin polymerization in vitro and for chloroplast movement in vivo (Figures 6, S6). These results demonstrate the structural and functional similarities between the CHUP1 C-terminal actin polymerization domain (CHUP1_716-982) and the FH2 domain of formin.

Formins are a major group of well-characterized actin nucleators that form tethered dimer structures (Xu et al., 2004) and are present in plants as well as other organisms (van Gisbergen and Bezanilla, 2013). Formins are involved in actin polymerization and associate with the fast-growing barbed ends of actin filaments. The proline-rich FH1 domain of formin mediates interactions with actin–profilin complexes, and the FH2 domain mediates actin polymerization by incorporating G-actin liberated from the actin–profilin complexes. The lasso domain of FH2 is important for its ability to form dimers. CHUP1-C forms dimers in solution (Figures 6B, 6C) that resemble FH2 dimers (Figures 4E, 4F) and which catalyze actin polymerization in the presence of profilin (Figure 5). Some formins, including one in plants (Yi et al., 2005), also have actin filament severing activity (Chhabra and Higgs, 2006; Harris et al., 2004). We showed by electron microscopy (Figures 5E-G, S3, S5) and fluorescence microscopy (Figure S2B) that CHUP1-C has actin filament severing activity in vitro, although we do not know whether CHUP1 severs actin filaments in vivo. These results suggest that CHUP1-C has structural homology to both the FH1 and FH2 domains of formin and has actin polymerization activity in the presence of profilin.

Together, our structural and biochemical analyses of the conserved CHUP1 C-terminal domain and our cell biological analyses of CHUP1 revealed that CHUP1 represents a novel class of plant-specific actin nucleator involved in chloroplast movement. Thus, CHUP1 is the fourth type of actin nucleator to be identified, joining the Arp2/3 complex (Goley and Welch, 2006), the formins (Chalkia et al., 2008), and Spire (Firat-Karalar and Welch, 2011).

### CHUP1 and cp-actin filaments mediate chloroplast movements that are important for plant survival

The chloroplast accumulation response is crucial for efficient photosynthesis and biomass production when light is limited (Gotoh et al., 2018), and the avoidance response minimizes photo-damage of chloroplasts in excess light (Kasahara et al., 2002). CHUP1 and cp-actin filaments are indispensable for these chloroplast movements. The distribution of CHUP1 on the chloroplast envelope varies, with different patterns, including spots, a diffuse distribution, and CHUP1 bodies, under different light conditions (Figures 1A, 7A). The dynamic reorganization of CHUP1 and of cp-actin filaments is reversibly regulated—mainly by phot2 but partially by phot1—in response to blue light (Figures 3E, 3F, 7B). During chloroplast photorelocation movements, cp-actin filaments are generated at the leading edge of each chloroplast, at the interface between the plasma membrane and the chloroplast (Kadota et al., 2009; Kong et al., 2013a). This asymmetric distribution of cp-actin filaments was coordinately regulated with the asymmetric distribution of CHUP1 on the front and rear ends of moving chloroplasts (Figures 3E, 3F, 7B). The reorganization of CHUP1 from dots into a diffuse pattern in the rear part of a moving chloroplast is probably related to cp-actin depolymerization by strong light. By contrast, small CHUP1 dots at the front of moving chloroplasts are probably involved in cp-actin polymerization. Therefore, CHUP1 determines the sites of cp-actin nucleation and polymerizes cp-actin filaments according to the intensity and position of incident blue light.

**Figure 7.**
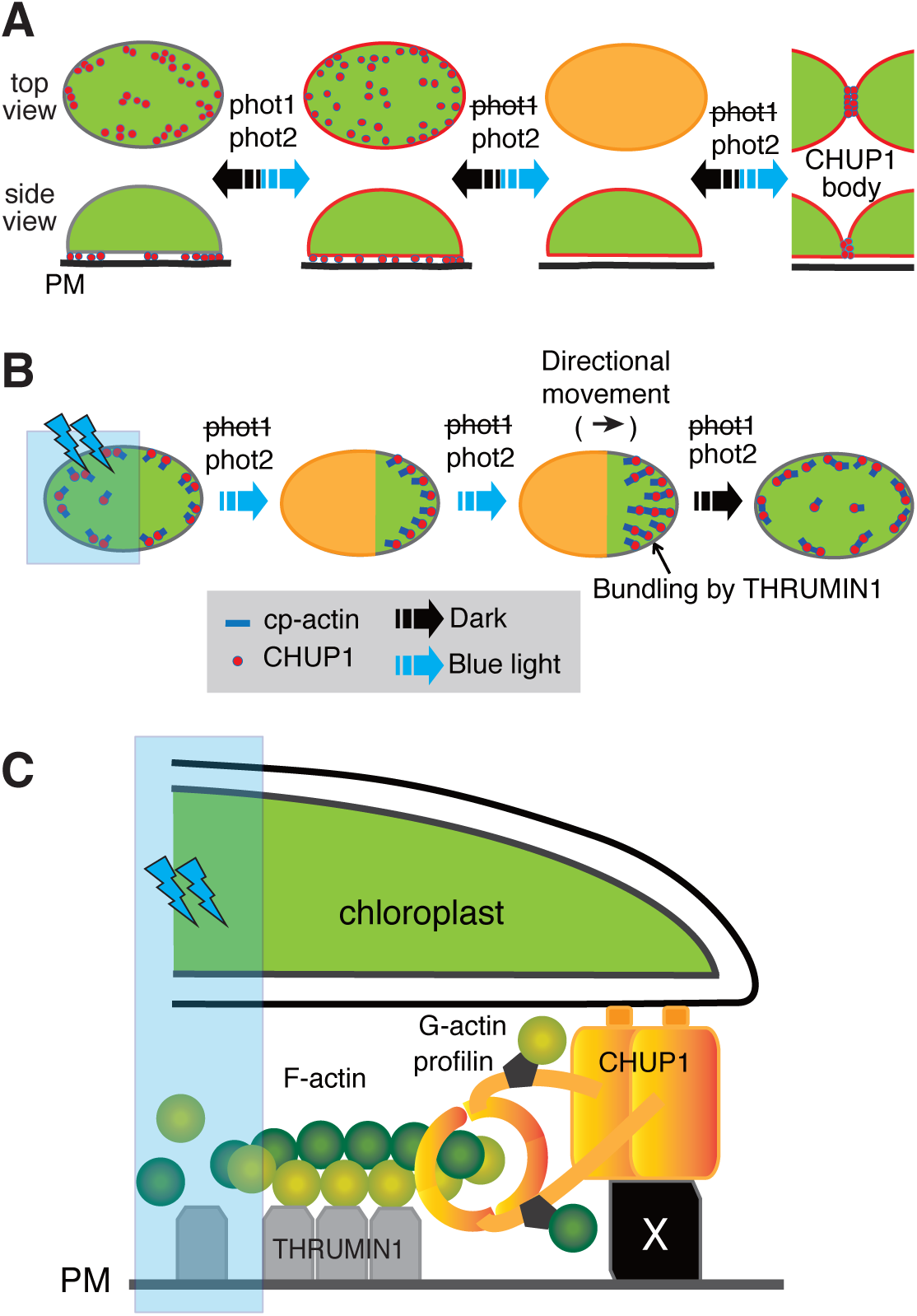
Schematic representation of coordinated phototropin-dependent dynamics of CHUP1 and cp-actin filaments. (A) Phototropin-dependent reorganization of CHUP1 (red) on the chloroplast envelope. CHUP1 localizes as dots at the interface between the plasma membrane and a chloroplast in darkness. In continuous strong light, CHUP1 disperses to the entire chloroplast envelope, first as dots at the interface and then diffusely throughout the chloroplast. Finally, CHUP1 accumulates as a CHUP1 body where two chloroplasts are in contact. These processes are reversibly regulated by phot2, but only partially by phot1. The lower panel shows side views of a chloroplast (side view) and the upper panel shows the chloroplast surface that faces the plasma membrane (top view). Blue arrows indicate phototropin-dependent processes under blue light and black arrows indicate the reverse processes in the dark. (B) Coordinated phototropin-dependent dynamics of CHUP1 and cp-actin filaments. Clusters of CHUP1 (red dots) nucleate cp-actin filaments. Asymmetric redistribution of CHUP1 and cp-actin filaments is coordinately regulated by phototropin in response to irradiation of part of a chloroplast with strong blue light (blue rectangle). phot2 is the main photoreceptor regulating the entire process of the strong light-induced avoidance response. (C) Working model of cp-actin-based chloroplast photorelocation. CHUP1 localizes to the chloroplast outer envelope via its N-terminal hydrophobic region. The coiled-coil domain of CHUP1 associates with an unidentified plasma membrane (PM)-localized protein (X), anchoring the chloroplast to the plasma membrane. A profilin-actin complex binds to the FH1-like domain of CHUP1-C and then the FH2-like domain of CHUP1-C adds G-actin to the barbed end of a cp-actin filament. The cp-actin filaments are bundled by PM-localized THRUMIN1. Eventually, the chloroplast is pushed forward by newly incorporated G-actin.

We do not know why plants evolved to have CHUP1 as a separate type of actin polymerization factor, when they also have their own formin family proteins. Interestingly, we identified sequences similar to the CHUP1 C-terminal half without the CHUP1 N-terminal half (named CHUC) in some non-Charophyte single-celled algae, including *Coccomyxa subellipsoidea* (Chlorophyta), *Micromonas commoda* (Chlorophyta), and *Guillardia theta* (Cryptophyta) as well as in land plants (Figure S7). These algae swim and their cell structures are compact, so chloroplast movement within the cell is not necessary. Thus, CHUC might not be involved in chloroplast movement in these algae. Consequently, the origin of CHUP1 and the physiological function of CHUC are unresolved.

### Does actin polymerization generate the motive force for chloroplast movement?

Some studies have indicated that the acto-myosin system is involved in chloroplast movement; that is, the chloroplasts move along a long cytoplasmic actin filament, driven by myosins. For example, a myosin XI tail domain was identified on the chloroplast envelope (Sattarzadeh et al., 2009), and the myosin inhibitors 2,3-butanedione monoxime (BDM) and N-ethylmaleimide (NEM) effectively inhibited chloroplast movement (Kong et al., 2013a; Paves and Truve, 2007; Yamada et al., 2011). However, double, triple, and quadruple knockout lines of the major myosin XIs in mesophyll cells of Arabidopsis leaves showed normal chloroplast movements, even though the movements of other organelles were defective (Suetsugu et al., 2010). We thus present an alternative model that relies on actin polymerization for generating motive force during chloroplast movements (Figure 7; Wada and Kong, 2018). In response to blue light, CHUP1 is reorganized and becomes asymmetrically redistributed to the front end of the chloroplast. Once that occurs, cp-actin filaments are polymerized at the interface between the plasma membrane and the chloroplast. Polymerized cp-actin filaments are bundled and anchored to the plasma membrane by a plasma membrane protein, THRUMIN1 (Whippo et al., 2011). Because the polymerized cp-actin filaments are anchored firmly to the plasma membrane, the cp-actin filaments that are newly polymerized by CHUP1 push the CHUP1 itself forward, thereby pushing the CHUP1-bound chloroplast forward. This proposed mechanism is reminiscent of the comet-tail movement of the intracellular bacterium *Listeria* in animal cells (Lambrechts et al., 2008). In the case of *Listeria*, actin polymerization is structurally restricted to one end of a rod-shaped bacterial cell, ensuring that the force vector of individual polymerizing actin filaments is oriented more or less parallel to the long axis of the cell. It remains to be elucidated how cp-actin filaments generate the motive force of chloroplast movement to ensure efficient photorelocation.

## STAR ★ METHODS

Detailed methods are provided in the online version of this paper and include the following:

- KEY RESOURCE TABLE
- CONTACT FOR REAGENT AND RESOURCE SHARING
- EXPERIMENTAL MODEL AND SUBJECT DETAILS
- METHOD DETAILS

## SUPPLEMENTAL INFORMATION

Supplemental Information includes nine figures, two tables, and seven movies and can be found with this article online at https://

## ACKNOWLEDGMENTS

The authors thank Takachika Tanaka, Yukiko Eda, and Hee-Jung Chung, for excellent technical assistance. We also thank the Arabidopsis Biological Resource Center for seed stocks. We thank the beamline staff at the SPring-8 beamlines—BL26B1, BL26B2, BL44XU, and BL32XU (Harima, Japan)—and the Photon Factory beamlines—BL-1A and BL-5A (Tsukuba, Japan)—for technical support. The experiments at the Photon Factory were performed with the approval of the Photon Factory Program Advisory Committee, as Proposal 2017G009, and those at SPring-8 BL44XU were performed under the Cooperative Research Program of the Institute for Protein Research, Osaka University, Osaka, Japan, as Proposals 20166617, 20176718, and 20186815. This work was supported in part by Grants-in-Aid for scientific research from the Ministry of Education, Culture, Sports, Science and Technology of Japan (MEXT)/the Japan Society for the Promotion of Science (JSPS) KAKENHI Grant Numbers 20227001, 23120523, 25120721, 25251033, and 16K14758 (to M.W); 21770050 (to S.-G.K.); 20687006, 24687014, and 25121726 (to A.S.); 26119002 (to D.K.); and 24117008 (to T.Q.P.U). Other support was provided by the Basic Science Research Program through the National Research Foundation of Korea (NRF), Ministry of Education (2016R1D1A3B03935947 to S.-G. K) and the Next-Generation Biogreen 21 Program grant, Rural Development Administration (RDA) (Project No. PJ01366901 to S.-G. K), and by the Ohsumi Frontier Science Foundation (to M.W.).

## AUTHOR CONTRIBUTIONS

S.-G.K., D.K., T.Q.P.U., and M.W. conceived this project. S.-G.K., A.S., D.K., T.Q.P.U., K.H., and M.W. planned the experiments and analyzed the data. S.-G.K., A.S., S.T.K., K.H., K.K., J.A., T.H., A.T., Y.N., N.S., and T.Q.P.U. carried out the experiments. S.-G.K., A.S., D.K., T.Q.P.U., and M.W. wrote the manuscript.

S.-G.K., K.K., J.A., T.H., N.S., and M.W. were involved in the plant physiological and cell biological experiments; A.S., A.T., Y.N., and D.K. in the crystallography; and S.-G.K., S.T.K., K.H., and T.Q.P.U. in the actin polymerization assays and EM observations.

### Accession Numbers

The coordinates and structure factors have been deposited in the Protein Data Bank, with the entries 6L2X for CHUP1_756-982 with the C864S mutation and 6L2Y for CHUP1_716-982, respectively.

## STAR ★ METHODS

### • KEY RESOURCES TABLE

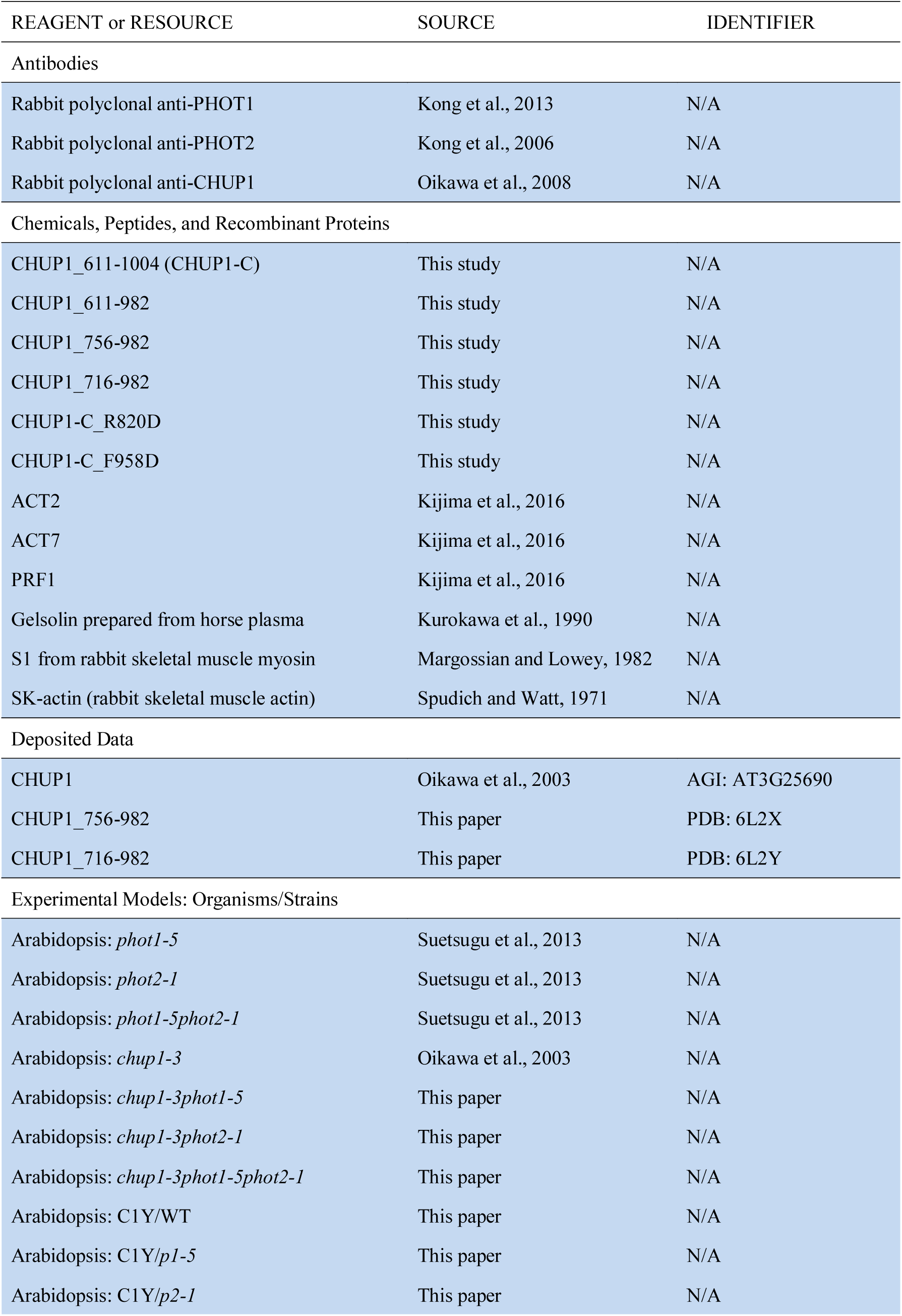
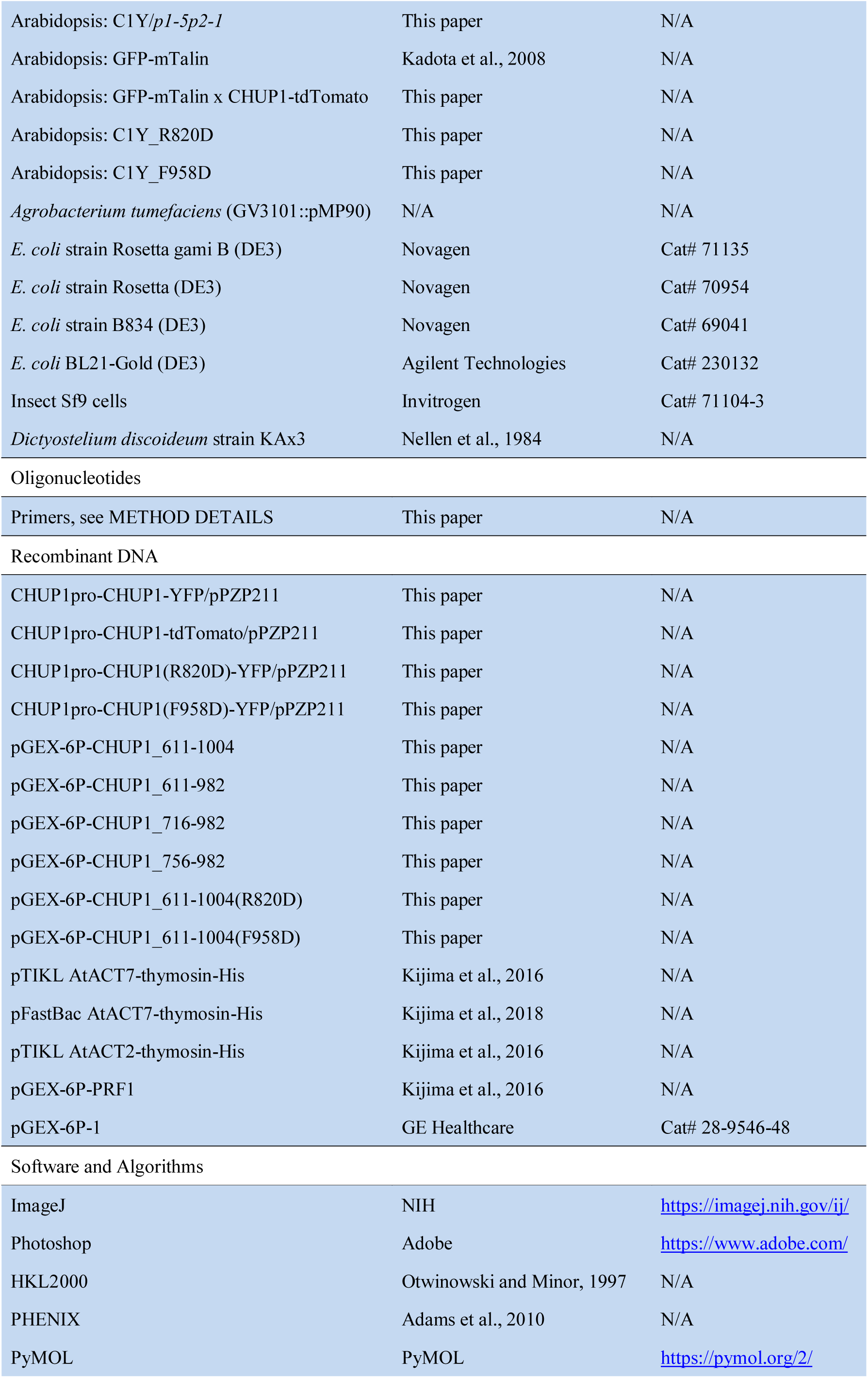
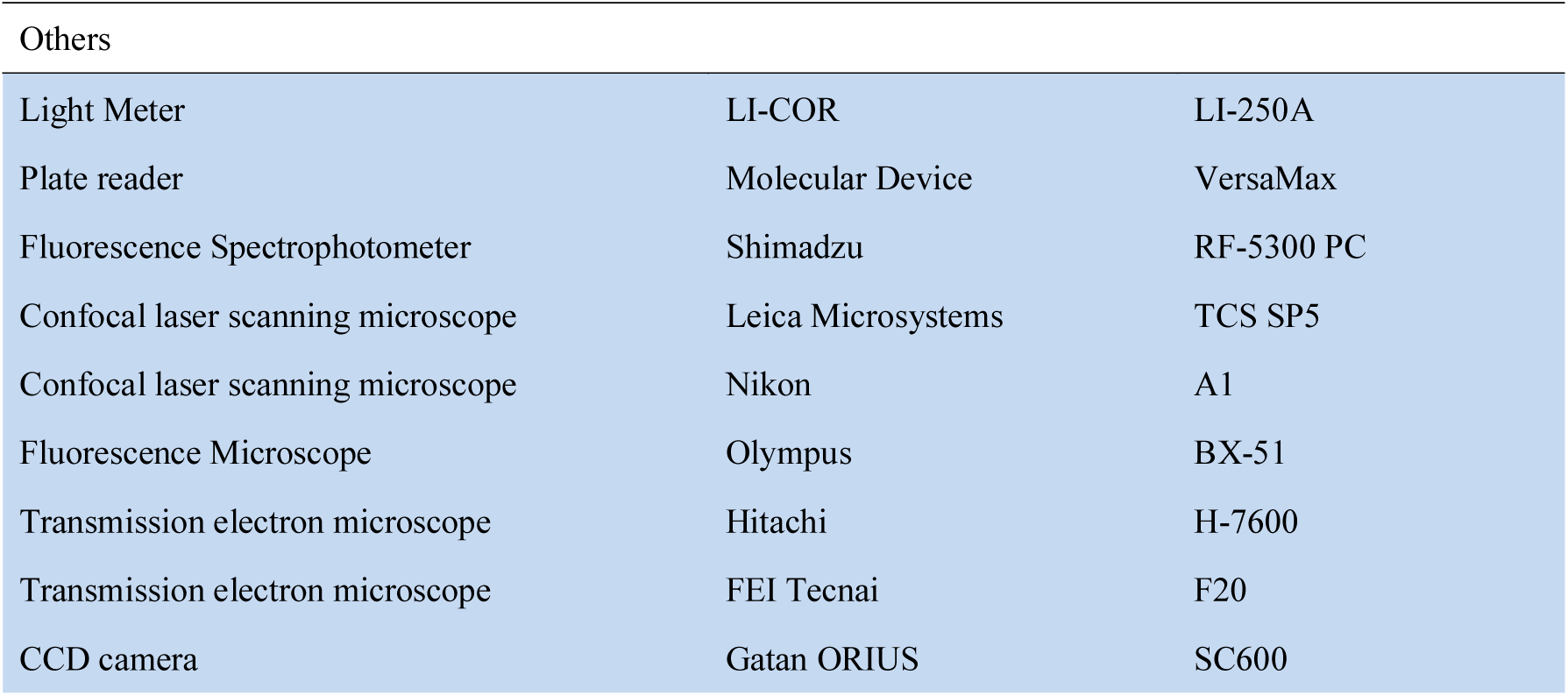

● KEY RESOURCES TABLE
| REAGENT or RESOURCE | SOURCE | IDENTIFIER |
| --- | --- | --- |
| Antibodies |  |  |
| Rabbit polyclonal anti-PHOT1 | Kong et al., 2013 | N/A |
| Rabbit polyclonal anti-PHOT2 | Kong et al., 2006 | N/A |
| Rabbit polyclonal anti-CHUP1 | Oikawa et al., 2008 | N/A |
| Chemicals, Peptides, and Recombinant Proteins |  |  |
| CHUP1_611-1004 (CHUP1-C) | This study | N/A |
| CHUP1_611-982 | This study | N/A |
| CHUP1_756-982 | This study | N/A |
| CHUP1_716-982 | This study | N/A |
| CHUP1-C_R820D | This study | N/A |
| CHUP1-C_F958D | This study | N/A |
| ACT2 | Kijima et al., 2016 | N/A |
| ACT7 | Kijima et al., 2016 | N/A |
| PRF1 | Kijima et al., 2016 | N/A |
| Gelsolin prepared from horse plasma | Kurokawa et al., 1990 | N/A |
| S1 from rabbit skeletal muscle myosin | Margossian and Lowey, 1982 | N/A |
| SK-actin (rabbit skeletal muscle actin) | Spudich and Watt, 1971 | N/A |
| Deposited Data |  |  |
| CHUP1 | Oikawa et al., 2003 | AGI: AT3G25690 |
| CHUP1_756-982 | This paper | PDB: 6L2X |
| CHUP1_716-982 | This paper | PDB: 6L2Y |
| Experimental Models: Organisms/Strains |  |  |
| Arabidopsis: *phot1-5* | Suetsugu et al., 2013 | N/A |
| Arabidopsis: *phot2-1* | Suetsugu et al., 2013 | N/A |
| Arabidopsis: *phot1-5phot2-1* | Suetsugu et al., 2013 | N/A |
| Arabidopsis: *chup1-3* | Oikawa et al., 2003 | N/A |
| Arabidopsis: *chup1-3phot1-5* | This paper | N/A |
| Arabidopsis: *chup1-3phot2-1* | This paper | N/A |
| Arabidopsis: *chup1-3phot1-5phot2-1* | This paper | N/A |
| Arabidopsis: C1Y/WT | This paper | N/A |
| Arabidopsis: C1Y/*p1-5* | This paper | N/A |
| Arabidopsis: C1Y/*p2-1* | This paper | N/A |
| Arabidopsis: C1Y/p1-5p2-1 | This paper | N/A |
| Arabidopsis: GFP-mTalin | Kadota et al., 2008 | N/A |
| Arabidopsis: GFP-mTalin x CHUP1-tdTomato | This paper | N/A |
| Arabidopsis: C1Y_R820D | This paper | N/A |
| Arabidopsis: C1Y_F958D | This paper | N/A |
| *Agrobacterium tumefaciens* (GV3101::pMP90) | N/A | N/A |
| *E. coli* strain Rosetta gami B (DE3) | Novagen | Cat# 71135 |
| *E. coli* strain Rosetta (DE3) | Novagen | Cat# 70954 |
| *E. coli* strain B834 (DE3) | Novagen | Cat# 69041 |
| *E. coli* BL21-Gold (DE3) | Agilent Technologies | Cat# 230132 |
| Insect Sf9 cells | Invitrogen | Cat# 71104-3 |
| *Dictyostelium discoideum* strain KAx3 | Nellen et al., 1984 | N/A |
| Oligonucleotides |  |  |
| Primers, see METHOD DETAILS | This paper | N/A |
| Recombinant DNA |  |  |
| CHUP1pro-CHUP1-YFP/pPZP211 | This paper | N/A |
| CHUP1pro-CHUP1-tdTomato/pPZP211 | This paper | N/A |
| CHUP1pro-CHUP1(R820D)-YFP/pPZP211 | This paper | N/A |
| CHUP1pro-CHUP1(F958D)-YFP/pPZP211 | This paper | N/A |
| pGEX-6P-CHUP1_611-1004 | This paper | N/A |
| pGEX-6P-CHUP1_611-982 | This paper | N/A |
| pGEX-6P-CHUP1_716-982 | This paper | N/A |
| pGEX-6P-CHUP1_756-982 | This paper | N/A |
| pGEX-6P-CHUP1_611-1004(R820D) | This paper | N/A |
| pGEX-6P-CHUP1_611-1004(F958D) | This paper | N/A |
| pTIKL AtACT7-thymosin-His | Kijima et al., 2016 | N/A |
| pFastBac AtACT7-thymosin-His | Kijima et al., 2018 | N/A |
| pTIKL AtACT2-thymosin-His | Kijima et al., 2016 | N/A |
| pGEX-6P-PRF1 | Kijima et al., 2016 | N/A |
| pGEX-6P-1 | GE Healthcare | Cat# 28-9546-48 |
| Software and Algorithms |  |  |
| ImageJ | NIH | https://imagej.nih.gov/ij/ |
| Photoshop | Adobe | https://www.adobe.com/ |
| HKL2000 | Otwinowski and Minor, 1997 | N/A |
| PHENIX | Adams et al., 2010 | N/A |
| PyMOL | PyMOL | https://pymol.org/2/ |
| Others |  |  |
| Light Meter | LI-COR | LI-250A |
| Plate reader | Molecular Device | VersaMax |
| Fluorescence Spectrophotometer | Shimadzu | RF-5300 PC |
| Confocal laser scanning microscope | Leica Microsystems | TCS SP5 |
| Confocal laser scanning microscope | Nikon | A1 |
| Fluorescence Microscope | Olympus | BX-51 |
| Transmission electron microscope | Hitachi | H-7600 |
| Transmission electron microscope | FEI Tecnai | F20 |
| CCD camera | Gatan ORIUS | SC600 |

### • CONTACT FOR REAGENT AND RESOURCE SHARING

Further information and requests for resources and reagents should be directed to and will be fulfilled by the Lead Contact, Masamitsu Wada

### • EXPERIMENTAL MODEL AND SUBJECT DETAILS

#### Plant materials

*Arabidopsis thaliana* ecotype Columbia was used for all experiments. The *chup1-3* mutant (Oikawa et al., 2003) was crossed with the *phot1-5*, *phot2-1*, and *phot1-5phot2-1* mutants (Suetsugu et al., 2013) to produce the mutants *chup1-3phot1-5*, *chup1-3phot2-1*, and *chup1-3phot1-5phot2-1*.

#### Transgenic plants

Transgenic Arabidopsis lines expressing CHUP1 fluorescent fusion proteins (CHUP1-tdTomato and CHUP1-YFP) from the native CHUP1 promoter were produced via *Agrobacterium*-mediated transformation of a CHUP1-deficient mutant (*chup1-3*) (Figure S1A). The CHUP1-YFP 10-5 transgenic line (C1Y/WT) was further crossed with the *chup1-3phot1-5*, *chup1-3phot2-1*, and *chup1-3phot1-5phot2-1* mutants to produce CHUP1pro::CHUP1-YFP/*phot1* (C1Y/*p1-5*), CHUP1pro::CHUP1-YFP/*phot2* (C1Y/*p2-1*), and CHUP1pro::CHUP1-YFP/*phot1phot2* (C1Y/*p1-5p2-1*) transgenic plants, respectively. The GFP-mTalin 2-28 transgenic line (Kadota et al., 2009; Kong et al., 2013a) was crossed with the CHUP1-tdTomato 8-1 transgenic line.

## METHOD DETAILS

### Immunoblot analysis

Arabidopsis plants were cultured on half-strength Murashige and Skoog (MS) medium (Nihon Pharmaceutical, Japan) supplemented with 1x MS vitamin (Sigma, USA), 1% (w/v) sucrose, and 0.8% (w/v) agar in a growth chamber set to a 16-h/8-h day/night cycle at 23°C. One hundred milligrams of rosette leaves frozen in liquid nitrogen was homogenized on ice using a glass homogenizer at a 1:3 (w:v) ratio of an extraction buffer (50 mM Tris-HCl, pH7.5, 10% (v/v) glycerol, 5 mM EDTA, 5 mM MgCl_2_, 1 mM dithiothreitol, and complete proteinase inhibitors) (Roche, Germany). The homogenates were clarified by centrifugation at 8,000 x *g* for 15 min at 4°C. Fifty micrograms of protein extracts was separated on 7.5% (w/v) SDS-PAGE gels, blotted onto a nitrocellulose membrane (Amersham, Switzerland), and probed with anti-CHUP1 (Oikawa et al., 2008), anti-phot1, or anti-phot2 (Kong et al., 2013b) polyclonal antibodies.

### Analysis of chloroplast photorelocation movement

Chloroplast movement was analyzed by measuring changes in red light transmittance, as previously described (Kong and Wada, 2011). For these assays, the fourth rosette leaves were detached from 3-week-old Arabidopsis plants cultured on half-strength MS medium. The detailed light conditions are described in the Figure 1 legend.

### Confocal microscopy

GFP, YFP, tdTomato, and chlorophyll fluorescence were detected using a Leica SP5 confocal microscope as previously described (Kong et al., 2013a). The excitation wavelength was 488 nm for GFP, 518 nm for YFP, and 561 nm for tdTomato. The emission ranges were 500–530 nm for GFP, 530–560 nm for YFP, and 575–630 nm for tdTomato.

The fluorescence images of GFP, tdTomato, and chlorophyll shown in Figure 3 were obtained using a Nikon A1 confocal microscope (Nikon, Tokyo) with a x 63 water immersion objective. The excitation lasers were 488 nm for GFP and 561 nm for tdTomato.

### Thin section electron microscopy

The detached fourth rosette leaves of 4-week-old WT plants were irradiated on agar plates with strong blue light (50 µmol m^−2^ s^−1^) for 30 min. The leaves were cut into about 2 mm x 2 mm pieces and were fixed in 2.5% (v/v) glutaraldehyde, postfixed in 1% (w/v) OsO_4_, dehydrated in acetone, and embedded in Spurr’s resin. Thin sections were stained with 4% (w/v) uranyl acetate and 0.4% (w/v) lead citrate and examined with a transmission electron microscope (model H-7600; Hitachi).

### Plasmid construction

cDNA fragments encoding the C-terminal fragments of CHUP1 including CHUP1_611-1104 (CHUP1-C), CHUP1_611-982, CHUP1_716-982, and CHUP1_756-982 were amplified from CHUP1 cDNA (AGI: AT3G25690) using PCR with the following set of primers.

The primer pair used to amplify the backbone of CHUP1_611-1004:

CHUP1_1831fw/*Bam*HI AA<u>GGATCC</u>ATCACAGCCACAGGTGATCAGT

CHUP1_3015rv(*)/*Sal*I AAGTCGACTCAGTTTACAGATTCTTCTTCATTG

The primer pair used to amplify the backbone of CHUP1_611-982:

CHUP1_1831fw/*Bam*HI AAGGATCCATCACAGCCACAGGTGATCAGT

CHUP1_2946rv(*)/*Sal*I AAGTCGACTCAGGCTCTGCTTCTAAGTTCCTC

The primer pair used to amplify the backbone of CHUP1_716-982:

CHUP1_2146fw/*Bam*HI AA<u>GGATCC</u>AACAAAGTTCACCGAGCTCCTGAG

CHUP1_2946rv(*)/*Sal*I AAGTCGACTCAGGCTCTGCTTCTAAGTTCCTC

The primer pair used to amplify the backbone of CHUP1_756-982:

CHUP1_2266fw/*Bam*HI AA<u>GGATCCAGGAACAATATGATTGGGGAAATC</u>

CHUP1_2946rv(*)/*Sal*I AAGTCGACTCAGGCTCTGCTTCTAAGTTCCTC

The PCR fragments were cloned into the *Bam*HI and *Sal*I restriction sites of the *Escherichia coli* expression vector pGEX-6P-1 (GE Healthcare) to produce pGEX-CHUP1 611-1004, pGEX-CHUP1_611-982, pGEX-CHUP1_716-982, and pGEX-CHUP1_756-982, respectively.

Genomic DNA from the promoter to the coding region of CHUP1 and the fluorescent genes of YFP, and tdTomato were amplified from genomic DNA prepared from Arabidopsis leaf tissues using PCR with the following primer pair.

CHUP1pro_(-2151)fw/*Hin*dIII AGTTGCTTAAAAGCTTGACTGAGGAGATTG

CHUP1_3012rv/*Sal*I AAGTCGACGTTTACAGATTCTTCTTCATTGCTA

The PCR fragment was cloned into the *Bam*HI and *Sal*I restriction sites of pPZP211/35S-nosT to produce CHUP1pro-CHUP1-YFP (or tdTomato)/pPZP211.

### Protein preparations for X-ray crystallography

Selenomethionine (SeMet)-substituted CHUP1_756-982 with a C864S mutation was expressed as a glutathione S-transferase (GST)-fusion protein using the pGEX-6P-1 vector (GE Healthcare). The transformed *Escherichia coli* strain B834(DE3) cells (Novagen) were grown at 37°C, until the OD_620_ reached 0.4 in the SeMet core medium (Wako), supplemented with 25 mg/L L-SeMet (Nacalai Tesque). After overnight induction with 0.5 mM isopropyl-β-D-thiogalactopyranoside (IPTG) at 20°C, the cells were harvested by centrifugation at 6,000 x *g* at 4°C for 15 min and resuspended in 50 mM Tris-HCl, pH 7.5, containing 500 mM NaCl, 2 mM 1,4-dithiothreithol (DTT), 2 mM EDTA, and 1% (v/v) glycerol (buffer A), supplemented with one tablet of cOmplete EDTA-free protease inhibitor (Roche) per 50 mL solution. After disruption by sonication and centrifugation at 18,000 x *g* at 4°C for 30 min, the supernatant was loaded onto a Glutathione Sepharose 4B (GE Healthcare) column equilibrated with buffer A. The column was then washed with buffer A. After cleavage with PreScission protease (GE Healthcare) on the column overnight, the protein was eluted with buffer A. The protein was further purified by gel filtration on a Superdex 200 10/300 GL column (GE Healthcare) in 50 mM Tris-HCl, pH 7.5, containing 200 mM NaCl and 2 mM DTT (buffer B). The purified protein was concentrated to 10 mg/mL for crystallization with an Amicon Ultra-15 centrifugal concentrator (Merck Millipore), and the buffer exchanged to 20 mM HEPES-NaOH, pH 7.0, containing 3 mM Tris(2-carboxyethyl)phosphine (TCEP) (buffer C).

The longer C-terminal fragment, CHUP1_716-982, was expressed as a GST-fusion protein using the pGEX-6P-1 vector. The transformed *E. coli* BL21-Gold (DE3) (Agilent Technologies) cells were grown at 37°C, until the OD_600_ reached 0.8. After overnight induction with 0.2 mM IPTG at 20°C, the cells were harvested by centrifugation at 6,000 x *g* at 4°C for 15 min and resuspended in 50 mM Tris-HCl, pH 8.0, containing 50 mM NaCl, 5 mM DTT, 2 mM MgCl_2_, and 10% (v/v) glycerol (buffer D), supplemented with one tablet of cOmplete EDTA-free protease inhibitor (Roche) and 5 µL Benzonase Nuclease (Merck Millipore) per 50 mL solution. After disruption by sonication and centrifugation at 18,000 x *g* at 4°C for 30 min, the supernatant was loaded on a Glutathione Sepharose 4B column equilibrated with buffer D. The column was sequentially washed with 50 mM Tris-HCl, pH 8.0, containing 400 mM NaCl, 5 mM DTT, and 10% (v/v) glycerol; 50 mM Tris-HCl, pH 8.0, containing 100 mM KCl, 10 mM MgCl_2_, 5 mM DTT, 0.25 mM ATP, and 5% (v/v) glycerol; and 50 mM Tris-HCl, pH 6.8, containing 150 mM NaCl, 1 mM EDTA (adjusted to pH 8.0), and 1 mM DTT (buffer E). After cleavage with PreScission protease on the column overnight, the protein was eluted with buffer E. The protein was further purified by gel filtration on a Superdex 200 10/300 GL column in 50 mM Tris-HCl, pH 8.0, containing 50 mM NaCl, 5 mM DTT, and 5% 32 (v/v) glycerol (buffer F). The purified protein was concentrated to 20 mg/mL with a Vivaspin Turbo 15 centrifugal concentrator (Sartorius), without changing the buffer composition.

### Crystallization and structure determination

CHUP1_756-982 and CHUP1_716-982 crystals were grown at 20°C using the hanging-drop vapor-diffusion method. The reservoir solution for CHUP1_756-982 was 50 mM CHES, pH 9.0, containing 0.2 M Li_2_SO_4_, and 36% PEG400, whereas that for CHUP1_716-982 was 1.1 M sodium malonate, 100 mM HEPES-NaOH, pH 7.4, containing 0.5% (v/v) Jeffamine ED-2001, pH 7.0 (Hampton Research). The crystals were briefly dipped into cryoprotectant solution containing 40% PEG400 for CHUP1_756-982 or 16% (v/v) glycerol for CHUP1_716-982, and then flash-cooled in liquid nitrogen.

Data sets were collected at the SPring-8 beamline BL44XU (Hyogo, Japan) and were processed with HKL2000 (Otwinowski and Minor, 1997). The structure of SeMet-substituted CHUP1_756-982 was solved by the SAD method, using the program PHENIX (Adams et al., 2010). The structure of the native CHUP1_716-982 was solved by the molecular replacement method with PHENIX, using the structure of the SeMet-substituted CHUP1_756-982 as a search model. Model building was refined using PHENIX. Figures were generated using PyMOL.

### Recombinant protein production for actin polymerization assay and EM observation

The proteins were expressed in *E. coli* strain Rosetta gami B (DE3) (Novagen) by 0.5 mM IPTG induction at 20°C after the *E. coli* cells were cultured to the late log phase (OD_600_ = 0.8 ∼1.0). The cells were harvested by centrifugation at 6,500 x *g* at 4°C for 20 min and homogenized in an extraction buffer (50 mM Tris-HCl, pH 7.5 at 23°C, 150 mM NaCl, 5% (v/v) glycerol, 5 mM DTT, 1% (v/v) Triton X-100, 1 mM EDTA, 0.2 mM PMSF). The GST proteins were trapped using a Glutathione Sepharose 4B matrix (GE Healthcare) and the CHUP1 C-terminal fragments were eluted by on-column cleavage using PreScission protease (GE Healthcare). The eluted CHUP1-C was further purified in a buffer (10 mM HEPES pH 7.4, 100 mM NaCl, 2 mM MgCl_2_, 1 mM DTT) by size-exclusion chromatography using a Superdex 200 10/300 column (GE Healthcare). The purified fractions were concentrated using a centrifugal filter unit (Amicon Ultra filter 30K, Millipore).

Arabidopsis profilin1 (PRF1) (AGI: AT2G19760) was expressed in *E. coli* strain Rosetta (DE3) (Novagen), and purified as described previously (Kijima et al., 2016). Arabidopsis vegetative actins ACT2 and ACT7 were expressed as fusion proteins with thymosin-His in Sf9 insect cells or *Dictyostelium discoideum*, and purified as described previously (Kijima et al., 2016). Rabbit skeletal muscle actin was prepared according to the method of Spudich and Watt (Spudich and Watt, 1971). Each purified protein was snap-frozen in liquid N_2_ and stored at –80°C.

### Actin Polymerization assay

Actin (4 µM) was allowed to polymerize in F-buffer (10 mM HEPES-NaOH pH 7.4, 50 mM KCl, 2 mM MgCl_2_, 0.2 mM ATP, 1 mM DTT) in the presence of various concentrations of PRF1 and CHUP1-C. After 30 min of incubation at 22°C, the solution was ultracentrifuged at 250,000 x *g* at 22°C for 10 min. The supernatant and pellet fractions were separately subjected to SDS-PAGE. The relative amounts of actin were estimated by analyzing intensities of Coomassie-stained bands using ImageJ (https://imagej.nih.gov/ij/)).

### Negative stain electron microscopy

Actin filaments diluted in EM buffer (10 mM potassium phosphate buffer pH 7.4, 25 mM KCl, 2.5 mM MgCl_2_, 0.2 mM ATP, and 0.5 mM DTT) were placed on carbon-coated copper grids, stained with 1% (w/v) uranyl acetate, and observed using an FEl Tecnai F20 electron microscope. The images were recorded with a Gatan ORIUS SC600 CCD camera, and processed with Adobe Photoshop to adjust the contrast and reduce noise.

## Supplemental Figures

**Figure S1. Complementation of chloroplast positioning and movements in C1Y transgenic plants**

(A) Schematic diagram of the CHUP1-YFP and -tdTomato expression vectors used for *Agrobacterium*-mediated transformation of *Arabidopsis thaliana*. The genomic DNA (pale green) for CHUP1pro::CHUP1-YFP (or -tdTomato; green) was cloned into the pPZP211/35S-nosT binary vector using the *Hin*dIII and *Pst*I restriction enzyme sites (Kong et al., 2013a). P_CHUP1_ indicates the native *CHUP1* promoter.

(B) Immunoblot analysis of CHUP1 and CHUP1-YFP in the rosette leaves of WT and C1Y transgenic plants, respectively. Fifty micrograms of total protein extracts were separated on 7.5% SDS-PAGE gels and probed with an anti-CHUP1 polyclonal antibody (Oikawa et al., 2008).

(C) Evaluation of chloroplast photorelocation movements in C1Y transgenic plants compared to those of WT and *chup1* mutant plants. The red-light transmittance in rosette leaves was monitored under three different intensities of blue light (3.2 µmol m^−2^ s^−1^, 25 µmol m^−2^ s^−1^, and 60 µmol m^−2^ s^−1^) for the indicated periods.

(D) Chloroplast distribution pattern in C1Y transgenic plants compared to those of WT and *chup1* mutant plants under different light conditions. The fourth rosette leaves of 4-week-old plants cultivated under a 16-h/ 8-h day/night cycle at 23°C were detached and incubated in darkness for 16 h and further under 2 µmol m^−2^ s^−1^ low-intensity blue light (LB) or 60 µmol m^−2^ s^−1^ high-intensity blue light (HB) for 2 h. The leaves were fixed with 2.5% glutaraldehyde in fixation solution (20 mM PIPES, 5 mM MgCl_2_, 5 mM EGTA, 0.5 mM phenylmethylsulfonyl fluoride, 1% dimethyl sulfoxide, pH7.0) for 30 min. The fixed leaves were subjected to confocal microscopy as previously described (Kong et al., 2007). Chloroplast position was visualized with red chlorophyll fluorescence. Scale bar = 50 µm.

**Figure S2. Interaction of CHUP1-C with skeletal muscle actin filaments**

(A) Interaction of CHUP1-C with the barbed actin filament ends. Rabbit skeletal muscle actin (SK-actin) was allowed to polymerize in F-buffer (10 mM HEPES pH 7.4, 100 mM NaCl, 2 mM MgCl_2_, 1 mM DTT and 0.1 mM ATP) containing 10 µM CaCl_2_ for 1 h. This was gently mixed with a solution of CHUP1-C diluted in F-buffer under various conditions (Sample 1–5), and after 10 min of incubation, the mixtures were ultracentrifuged at 250,000 x *g* for 10 min. The final concentrations of actin and CHUP1-C were 10 µM and 1 µM, respectively, and the temperature was 22°C throughout the procedure. The supernatant and pellet fractions were separately subjected to SDS-PAGE. Sample 1: control. Sample 2: actin filaments were sheared by vigorous pipetting (30 times, up and down) immediately before the addition of CHUP1-C. Sample 3: Actin filaments were fragmented by 0.1 µM gelsolin immediately before the addition of CHUP1-C. Plasma gelsolin was purified from horse plasma as described (Kurokawa et al., 1990). Sample 4: the same as for sample 1, except that 1 mM EGTA was added to chelate Ca^2+^ before the addition of CHUP1-C. Sample 5: CHUP1-C without actin filaments. The arrow and double arrow on the right indicate the positions of actin and CHUP1-C, respectively.

(B) Fluorescence images of actin filaments after staining with rhodamine-phalloidin and immobilization on S1-coated coverslips. From left to right, control, sheared by vigorous pipetting (without CHUP1-C), and supplemented with 1 µM CHUP1-C (the same as for sample 1 in A). Scale bar = 10 µm. The lengths of more than 200 actin filaments were measured for each condition, and the average and SD were calculated.

(C) Polymerization of skeletal muscle actin in the presence of various concentrations of CHUP1-C. Actin polymerization was monitored by measuring the increase in the fluorescence intensity of pyrene-labeled actin (Kouyama and Mihashi, 1981).

**Figure S3. Electron micrographs of negatively stained rabbit skeletal muscle actin filaments**

(A, D) Control rabbit skeletal muscle actin (SK-actin, 8 µM) polymerized at room temperature for 30 min in F-buffer.

(B, E) Actin filaments 15 s after the addition of 2 µM CHUP1-C.

(C, F) Actin filaments 15 min after the addition of 2 µM CHUP1-C.

Higher-magnification images (D–F) revealed that filaments incubated with CHUP1-C had locally thick and narrow sections or gaps. The narrow sections may represent gaps in one of the protofilaments (blue arrows in E and F). The thick sections may represent bound CHUP1-C along the sides, and notably, blob-like structures were often found juxtaposing the narrow sections (red arrows in E and F).

(G) Actin filaments polymerized at 8 µM in the presence of 2 µM CHUP1-C at room temperature for 45 min. Large structures were often observed at the filament ends (blue arrows).

**Figure S4. Kinetic analysis of ACT7 polymerization in the presence of PRF1 and CHUP1-C**

Salt-induced polymerization was tracked by measuring the fluorescence intensity of pyrene attached to skeletal muscle actin (0.4 µM) that was included as the probe in the reaction mixture. Graph A: increasing the concentration of PRF1 (from 0 to 8 µM) inhibited polymerization of 4 µM ACT7. Graph B: the inhibitory effect of PRF1 on polymerization of ACT7 was rescued by 1 µM CHUP1-C in a manner partially dependent on PRF1. The fluorescence intensities in the presence of PRF1 did not reach those in the absence of PRF1, even though ultracentrifugation assays (Figure 6D) demonstrated that most of the ACT7 polymerized under the same condition. A similar apparent discrepancy was reported when muscle actin, mDia1, and sheep spleen profilin were combined (Romero et al., 2004), and this was attributed to poor binding of pyrenyl actin to profilin.

**Figure S5. Electron micrographs of ACT7 filaments incubated with CHUP1-C**

(A) Control filaments without CHUP1-C.

(B–C) ACT7 filaments (0.56 µM) negatively stained immediately after the addition of 1 µM CHUP1-C. This is similar to the condition of Figure 5F, except that in the experiment shown in Figure 5F, ACT7 filaments and CHUP1-C were incubated in a tube for 1–2 minutes before being applied to a carbon-coated copper grid, whereas filaments shown here were applied to the grid and negatively stained within several seconds of CHUP1-C addition.

**Figure S6. Protein sequence alignment of the C-terminal region of CHUP1**

All aligned sequences were obtained from GenBank. Green bar, profilin-rich region of the FH1-like domain. Blue bar, FH2-like domains conserved in CHUP1 family proteins. Red boxes, R820 and F958.

Ac, *Adiantum capillus-veneris* (Ferns); At, *Arabidopsis thaliana* (Angiosperms); Atr, *Amborella trichopoda* (Angiosperms); Kf, *Klebsormidium flaccidum* (Klebsormidiales); Lj, *Lygodium japonicum* (Ferns); Mp, *Marchantia polymorpha* (Liverworts); Mv, *Mesostigma viride* (Mesostigmatales); Nn, *Nelumbo nucifera* (Angiosperms); Os, *Oryza sativa* (Angiosperms); Pp, *Physcomitrella patens* (Mosses); Sm, *Selaginella moellendorffii* (Lycophytes).

**Figure S7. Organismal lineages and distributions of CHUP1 and CHUC genes**

The topology of lineages is adapted from (Li et al., 2015). CHUP1 genes were identified in the genome and/or transcriptome databases of charophytes and land plants but not in those of chlorophytes. By contrast, CHUC genes were identified in unicellular algae such as *Guillardia theta*, *Micromonas commoda*, and *Coccomyxa subellipsoidea* as well as in land plants.

## Supplemental Tables

**Table S1.**
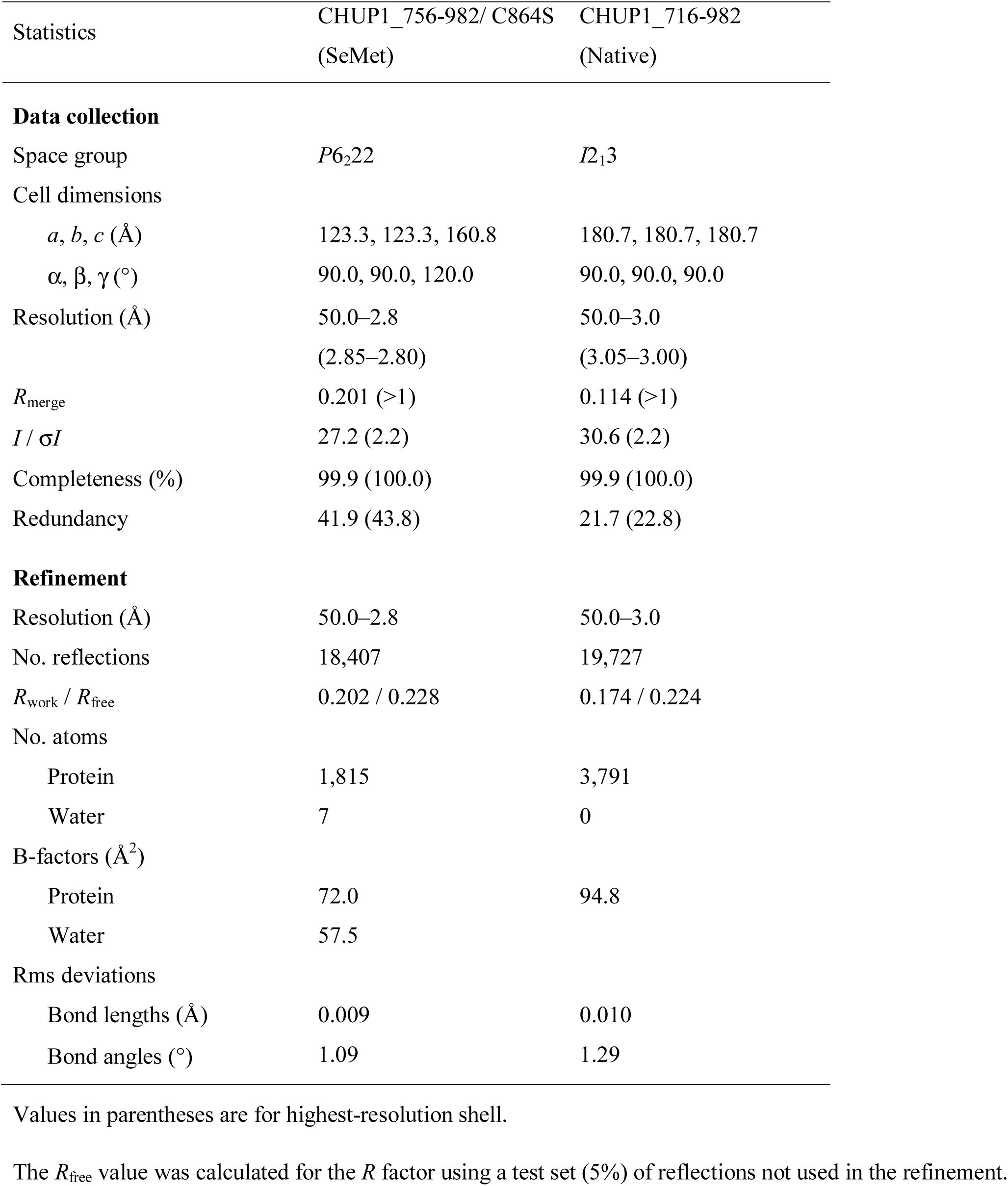
Data collection and refinement statistics.

**Table S2.**
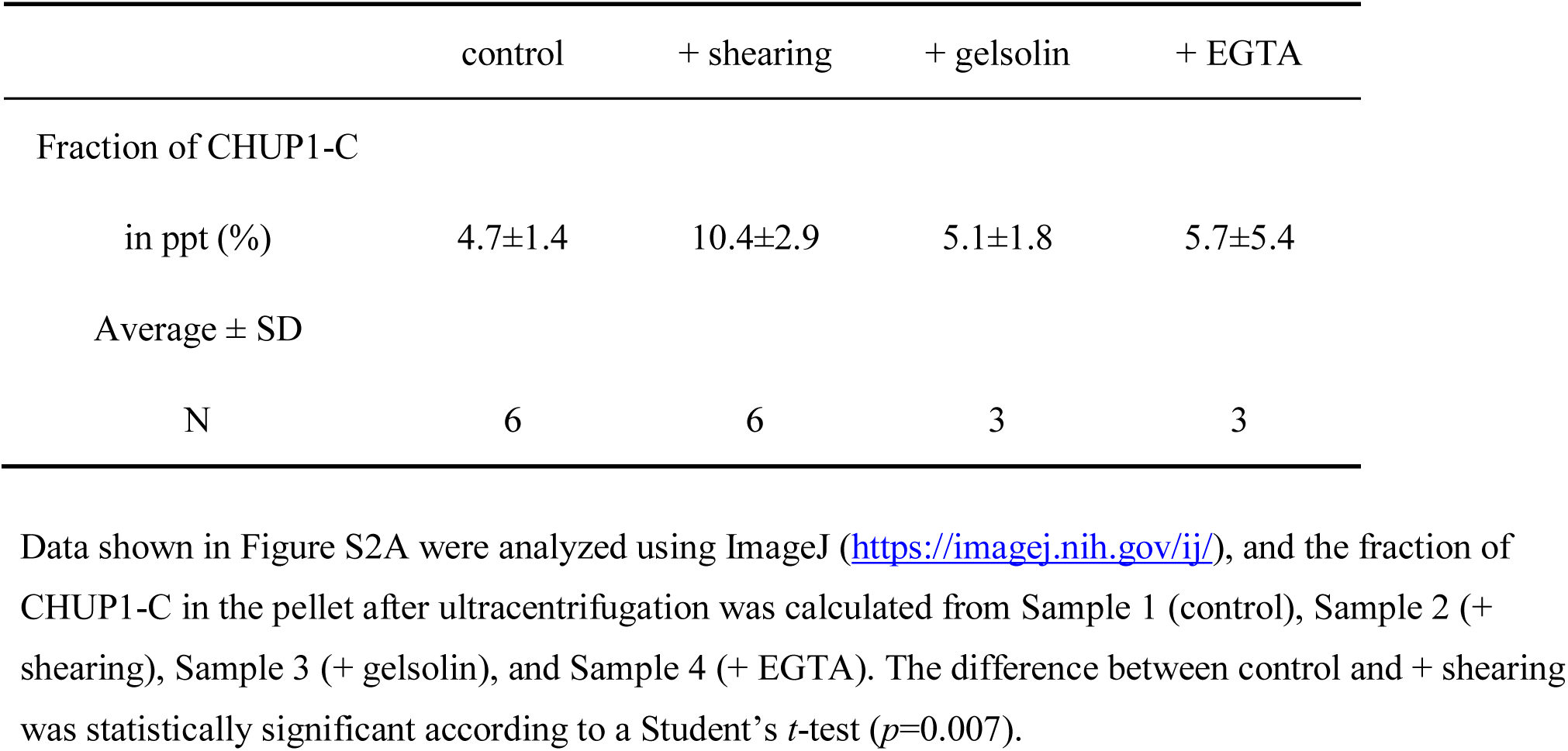
Binding of CHUP1-C to skeletal muscle actin filaments.

## Supplemental Movies

**Movie S1. Reorganization of CHUP1-YFP in a WT palisade cell during the blue light-induced chloroplast avoidance response**

Same cell as in Figure 1D. Time-lapse images (maximized with three images in a 1.2-µm depth) were collected at approximately 40-s intervals and played back at 5 frames per second (fps). The total elapsed time was 13:30 (min:s). The images are false-colored to indicate YFP (yellow) and chlorophyll (red) fluorescence. The region indicated by the blue circle was irradiated using 458-nm laser scans in the intervals between image acquisitions. Scale bar = 10 µm.

**Movie S2. Reorganization of CHUP1-YFP in a WT palisade cell during the blue light-induced chloroplast avoidance response**

Same cell as in Figure 1G. The total elapsed time is 10:30 (min:s). Other details are the same as in Movie S1. Note that the chloroplast in the roi cannot avoid strong light without resulting in the asymmetric distribution of CHUP1-YFP.

**Movie S3. Reorganization of CHUP1-Y in a WT palisade cell during the blue light-induced chloroplast avoidance response**

Same cell as in Figure 2B. The total elapsed time is 10:27 (min:s). The region indicated with the blue rectangle was irradiated by 458-nm laser scans. Scale bar = 5 µm. Other details are the same as in Movie S1.

**Movie S4. Reorganization of CHUP1-YFP in a *phot1* mutant palisade cell during the blue light-induced chloroplast avoidance response.**

The same cell as in Figure 2B. The total elapsed time was 10:27 (min:s). Other details are the same as in Movie S3.

**Movie S5. Reorganization of CHUP1-YFP in a *phot2* mutant palisade cell during the blue light-induced chloroplast avoidance response**

The same cell as in Figure 2B is shown. The total elapsed time was 10:00 (min:s). Other details are the same as in Movie S3.

**Movie S6. Reorganization of CHUP1-YFP in a *phot1phot2* mutant palisade cell during the blue light-induced chloroplast avoidance response**

The same cell as in Figure 2B is shown. The total elapsed time was 10:30 (min:s). Other details are the same as in Movie S3.

**Movie S7. Reorganizations of CHUP1 and cp-actin filaments in a CHUP1-tdTomato x GFP-mTalin palisade cell during the blue light-induced chloroplast avoidance response**

Same cell as in Figure S3F. Time-lapse images were collected at approximately 34-s intervals. The total elapsed time was 10:55 (min:s). The images were false-colored to indicate GFP (green), tdTomato (red), and chlorophyll (gray) fluorescence. The region indicated by the blue rectangle (15 µm x 20 µm) was irradiated. Scale bar = 10 µm. Other details are the same as in Movie S1.

