## Supplementary Data for "CHLOROPLAST UNUSUAL POSITIONING 1 is a new type of actin nucleation factor in plants"

### Supplemental Figures

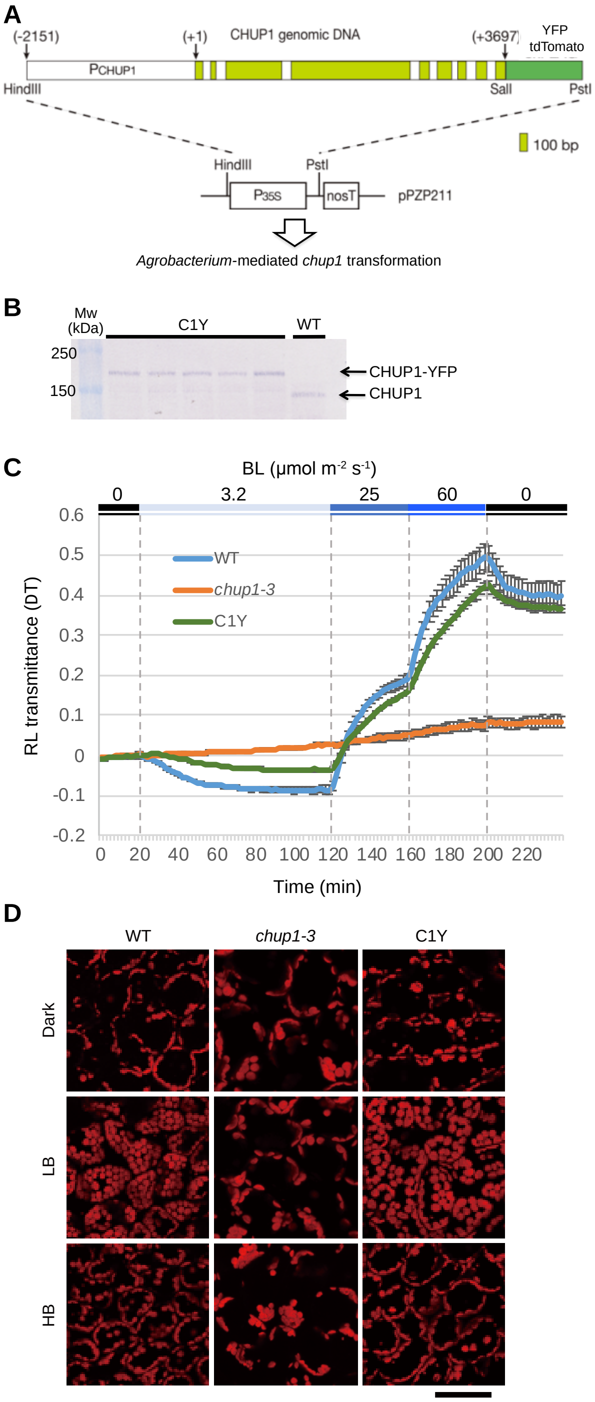


**Figure S1. Complementation of chloroplast positioning and movements in C1Y transgenic plants**

(A) Schematic diagram of the CHUP1-YFP and -tdTomato expression vectors used for *Agrobacterium*-mediated transformation of *Arabidopsis thaliana*. The genomic DNA (pale green) for

CHUP1pro::CHUP1-YFP (or -tdTomato; green) was cloned into the pPZP211/35S-nosT binary vector using the *Hin*dIII and *Pst*I restriction enzyme sites (Kong et al., 2013a). $P_{CHUP1}$ indicates the native *CHUP1* promoter.

(B) Immunoblot analysis of CHUP1 and CHUP1-YFP in the rosette leaves of WT and C1Y transgenic plants, respectively. Fifty micrograms of total protein extracts were separated on 7.5% SDS-PAGE gels and probed with an anti-CHUP1 polyclonal antibody (Oikawa et al., 2008).

(C) Evaluation of chloroplast photorelocation movements in C1Y transgenic plants compared to those of WT and *chup1* mutant plants. The red-light transmittance in rosette leaves was monitored under three different intensities of blue light (3.2 µmol $m^{-2}$ $s^{-1}$, 25 µmol $m^{-2}$ $s^{-1}$, and 60 µmol $m^{-2}$ $s^{-1}$) for the indicated periods.

(D) Chloroplast distribution pattern in C1Y transgenic plants compared to those of WT and *chup1* mutant plants under different light conditions. The fourth rosette leaves of 4-week-old plants cultivated under a 16-h/ 8-h day/night cycle at 23°C were detached and incubated in darkness for 16 h and further under 2 µmol $m^{-2}$ $s^{-1}$ low-intensity blue light (LB) or 60 µmol $m^{-2}$ $s^{-1}$ high-intensity blue light (HB) for 2 h. The leaves were fixed with 2.5% glutaraldehyde in fixation solution (20 mM PIPES, 5 mM $MgCl_2$, 5 mM EGTA, 0.5 mM phenylmethylsulfonyl fluoride, 1% dimethyl sulfoxide, pH7.0) for 30 min. The fixed leaves were subjected to confocal microscopy as previously described (Kong et al., 2007). Chloroplast position was visualized with red chlorophyll fluorescence. Scale bar = 50 µm.

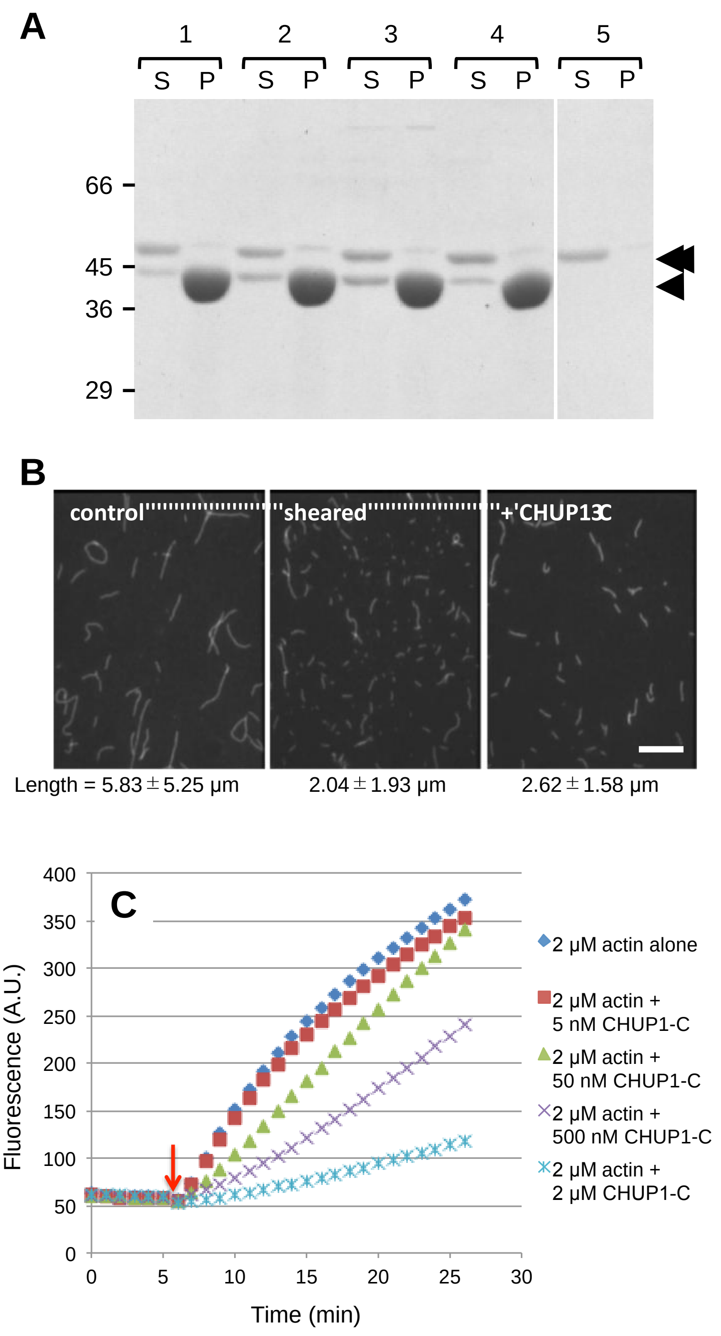


**Figure S2. Interaction of CHUP1-C with skeletal muscle actin filaments**

(A) Interaction of CHUP1-C with the barbed actin filament ends. Rabbit skeletal muscle actin (SK-actin) was allowed to polymerize in F-buffer (10 mM HEPES pH 7.4, 100 mM NaCl, 2 mM $MgCl_2$, 1 mM DTT and 0.1 mM ATP) containing 10 μM $CaCl_2$ for 1 h. This was gently mixed with a solution of CHUP1-C diluted in F-buffer under various conditions (Sample 1–5), and after 10 min of incubation, the mixtures were ultracentrifuged at 250,000 x *g* for 10 min. The final concentrations of actin and CHUP1-C were 10 μM and 1 μM, respectively, and the temperature was 22°C throughout the procedure. The supernatant and pellet fractions were separately subjected to SDS-PAGE. Sample 1: control. Sample 2: actin filaments were sheared by vigorous pipetting (30 times, up and down) immediately before the addition of CHUP1-C. Sample 3: Actin filaments were fragmented by 0.1 μM gelsolin immediately before the addition of CHUP1-C. Plasma gelsolin was purified from horse plasma as described (Kurokawa et al., 1990). Sample 4: the same as for sample 1, except that 1 mM EGTA was added to chelate $Ca^{2+}$ before the addition of CHUP1-C. Sample 5: CHUP1-C without actin filaments. The arrow and double arrow on the right indicate the positions of actin and CHUP1-C, respectively.

(B) Fluorescence images of actin filaments after staining with rhodamine-phalloidin and immobilization on S1-coated coverslips. From left to right, control, sheared by vigorous pipetting (without CHUP1-C), and supplemented with 1 μM CHUP1-C (the same as for sample 1 in A). Scale bar = 10 μm. The lengths of more than 200 actin filaments were measured for each condition, and the average and SD were calculated.

(C) Polymerization of skeletal muscle actin in the presence of various concentrations of CHUP1-C. Actin polymerization was monitored by measuring the increase in the fluorescence intensity of pyrene-labeled actin (Kouyama and Mihashi, 1981).

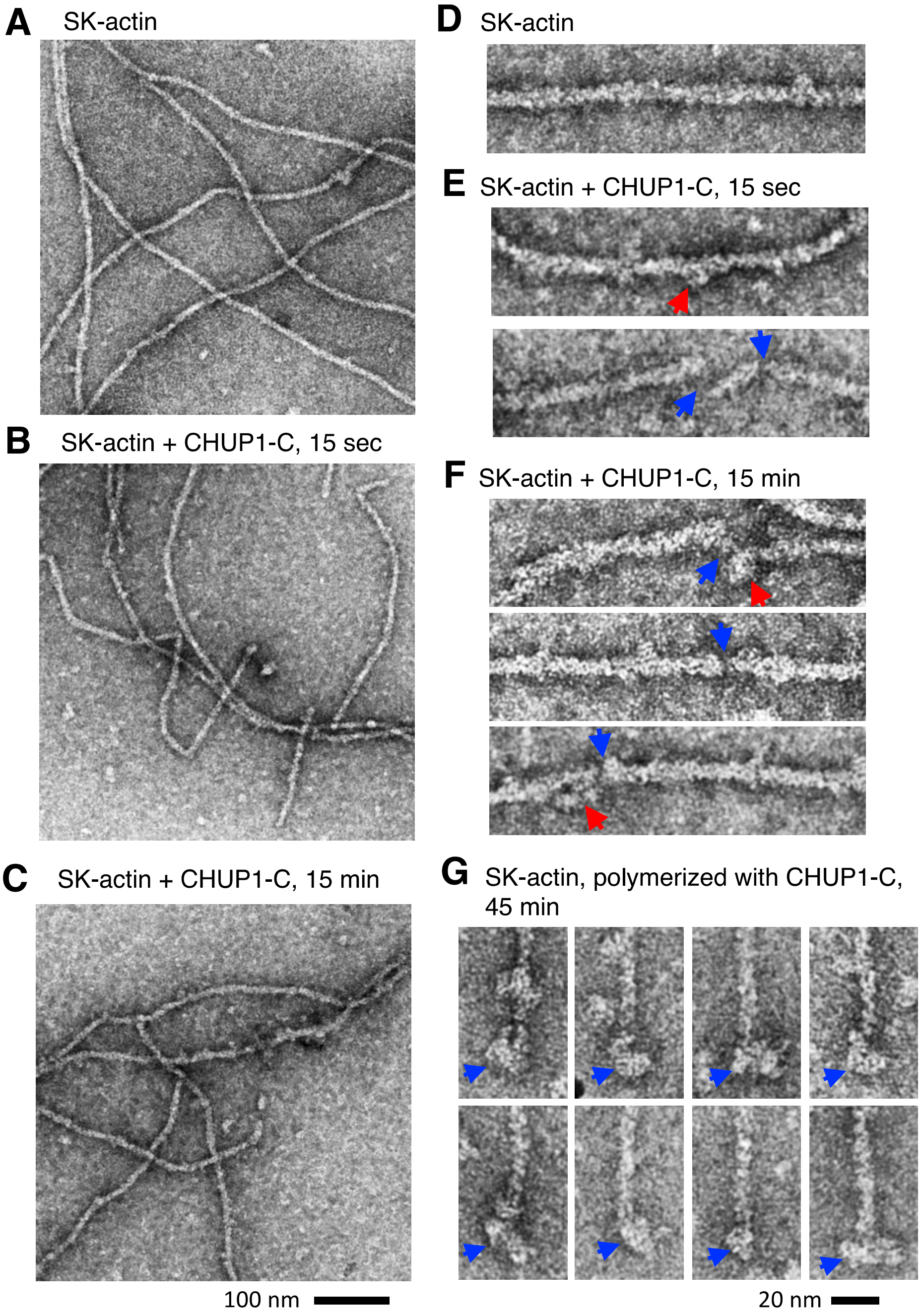


**Figure S3. Electron micrographs of negatively stained rabbit skeletal muscle actin filaments**

(A, D) Control rabbit skeletal muscle actin (SK-actin, 8 μM) polymerized at room temperature for 30 min in F-buffer.

(B, E) Actin filaments 15 s after the addition of 2 μM CHUP1-C.

(C, F) Actin filaments 15 min after the addition of 2 μM CHUP1-C.

Higher-magnification images (D–F) revealed that filaments incubated with CHUP1-C had locally thick and narrow sections or gaps. The narrow sections may represent gaps in one of the protofilaments (blue arrows in E and F). The thick sections may represent bound CHUP1-C along the sides, and notably, blob-like structures were often found juxtaposing the narrow sections (red arrows in E and F).

(G) Actin filaments polymerized at 8 μM in the presence of 2 μM CHUP1-C at room temperature for 45 min. Large structures were often observed at the filament ends (blue arrows).

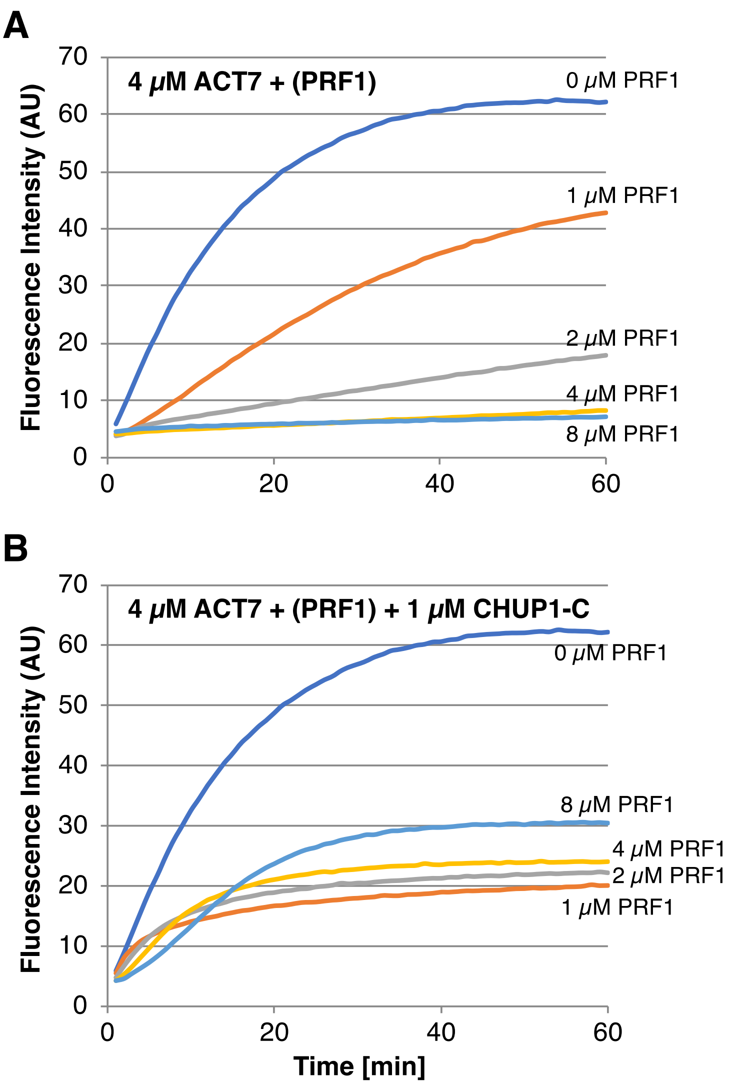


**Figure S4. Kinetic analysis of ACT7 polymerization in the presence of PRF1 and CHUP1-C**

Salt-induced polymerization was tracked by measuring the fluorescence intensity of pyrene attached to skeletal muscle actin (0.4 μM) that was included as the probe in the reaction mixture. Graph A: increasing the concentration of PRF1 (from 0 to 8 μM) inhibited polymerization of 4 μM ACT7. Graph B: the inhibitory effect of PRF1 on polymerization of ACT7 was rescued by 1 μM CHUP1-C in a manner partially dependent on PRF1. The fluorescence intensities in the presence of PRF1 did not reach those in the absence of PRF1, even though ultracentrifugation assays (Figure 6D) demonstrated that most of the ACT7 polymerized under the same condition. A similar apparent discrepancy was reported when muscle actin, mDia1, and sheep spleen profilin were combined (Romero et al., 2004), and this was attributed to poor binding of pyrenyl actin to profilin.

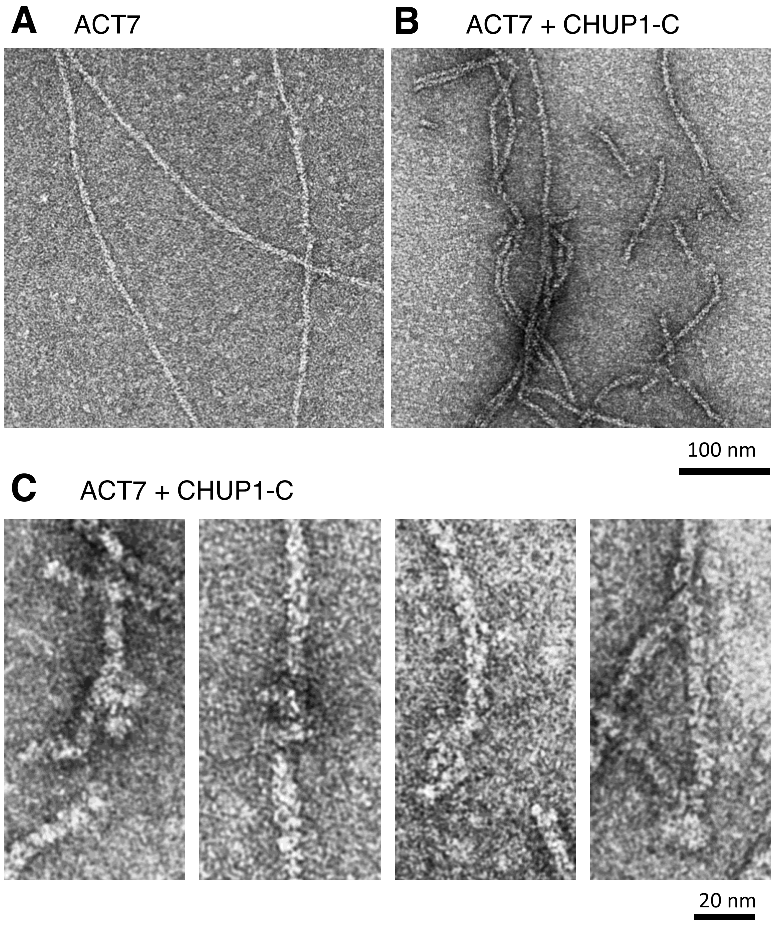


**Figure S5. Electron micrographs of ACT7 filaments incubated with CHUP1-C**

(A) Control filaments without CHUP1-C.

(B–C) ACT7 filaments (0.56 µM) negatively stained immediately after the addition of 1 µM CHUP1-C. This is similar to the condition of Figure 5F, except that in the experiment shown in Figure 5F, ACT7 filaments and CHUP1-C were incubated in a tube for 1–2 minutes before being applied to a carbon-coated copper grid, whereas filaments shown here were applied to the grid and negatively stained within several seconds of CHUP1-C addition.

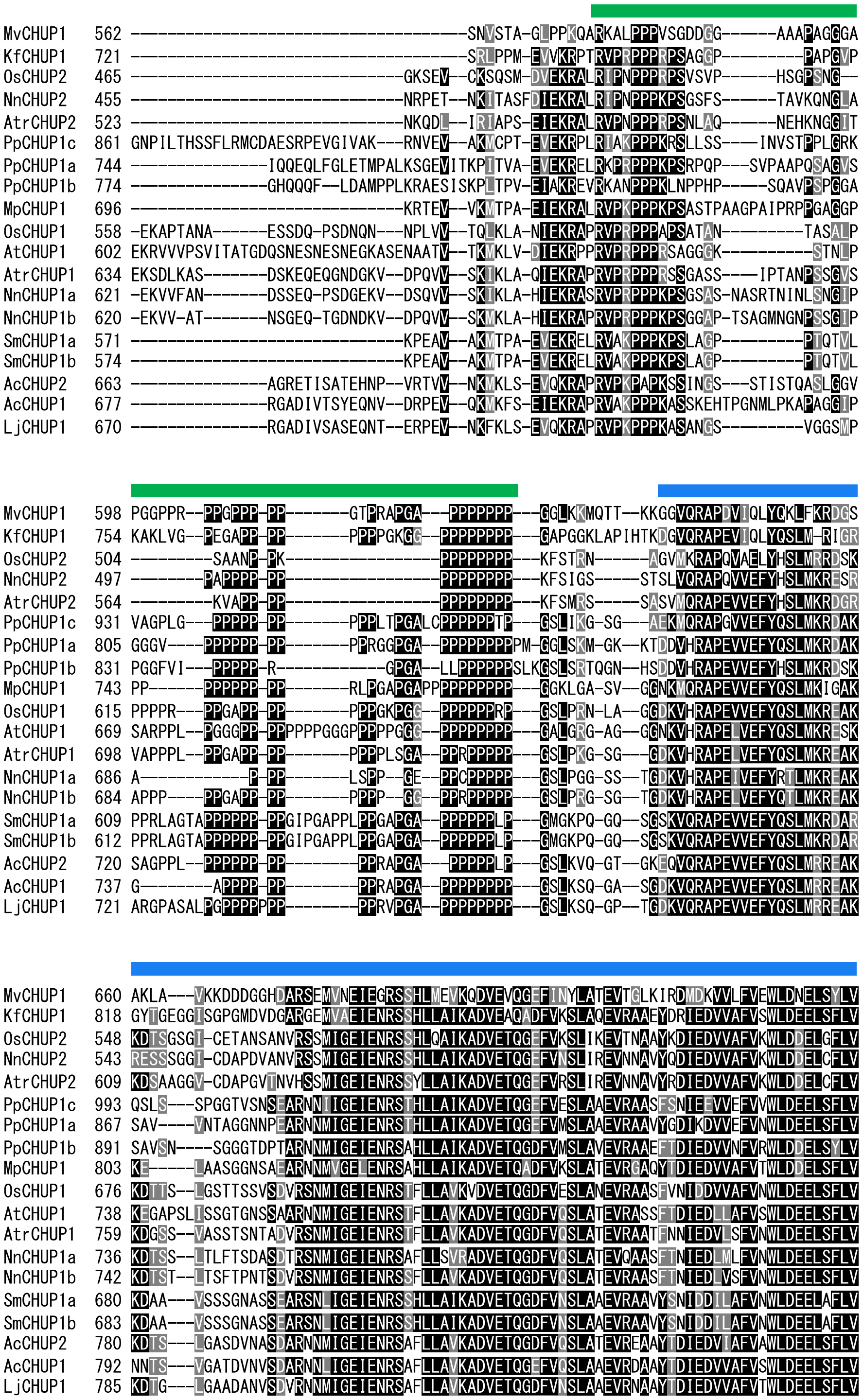
MvCHUP1 562 -------------------------------------SNVSTA-GLPPKQARKALPPPVSGDDGG------AAAPAGGGA
KfCHUP1 721 -------------------------------------SRLPPM-EVVKRPTRVPRPPPRPSAGGP---------PAPGVP
OsCHUP2 465 ------------------------------GKSEV--CKSQSM-DVEKRALRIPNPPPRPSVSVP------HSGPSNG--
NnCHUP2 455 ------------------------------NRPET--NKITASFDIEKRALRIPNPPPKPSGSFS------TAVKQNGLA
AtrCHUP2 523 ------------------------------NKQDL--IRIAPS-EIEKRALRVPNPPPRPSNLAQ------NEHKNGGIT
PpCHUP1c 861 GNPILTHSSFLRMCDAESRPEVGIVAK---RNVEV--AKMCPT-EVEKRPLRIAKPPPKRSLLSS----INVSTPPLGRK
PpCHUP1a 744 ---------------IQQEQLFGLETMPALKSGEVITKPITVA-EVEKRELRKPRPPPKPSRPQP---SVPAAPQSAGVS
PpCHUP1b 774 ---------------GHQQQF--LDAMPPLKRAESISKPLTPV-EIAKREVRKANPPPKLNPPHP-----SQAVPSPGGA
MpCHUP1 696 ------------------------------KRTEV--VKMTPA-EIEKRALRVPKPPPKPSASTPAAGPAIPRPPGAGGP
OsCHUP1 558 -EKAPTANA------ESSDQ-PSDNQN---NPLVV--TQLKLA-NIEKRAPRVPRPPPAPSATAN---------TASALP
AtCHUP1 602 EKRVVVPSVITATGDQSNESNESNEGKASENAATV--TKMKLV-DIEKRPPRVPRPPPRSAGGGK----------STNLP
AtrCHUP1 634 EKSDLKAS-------DSKEQEQGNDGKV--DPQVV--SKIKLA-QIEKRAPRVPRPPPRSSGASS----IPTANPSSGVS
NnCHUP1a 621 -EKVVFAN-------DSSEQ-PSDGEKV--DSQVV--SKIKLA-HIEKRASRVPRPPPKPSGSAS-NASRTNINLSNGIP
NnCHUP1b 620 -EKVV-AT-------NSGEQ-TGDNDKV--DPQVV--SKMKLA-HIEKRAPRVPRPPPKPSGGAP-TSAGMNGNPSSGIP
SmCHUP1a 571 ------------------------------KPEAV--AKMTPA-EVEKRELRVAKPPPKPSLAGP---------PTQTVL
SmCHUP1b 574 ------------------------------KPEAV--AKMTPA-EVEKRELRVAKPPPKPSLAGP---------PTQTVL
AcCHUP2 663 ---------------AGRETISATEHNP--VRTVV--NKMKLS-EVQKRAPRVPKPAPKSSINGS---STISTQASLGGV
AcCHUP1 677 ---------------RGADIVTSYEQNV--DRPEV--QKMKFS-EIEKRAPRVAKPPPKASSKEHTPGNMLPKAPAGGIP
LjCHUP1 670 ---------------RGADIVSASEQNT--ERPEV--NKFKLS-EVQKRAPRVPKPPPKASANGS---------VGGSMP
MvCHUP1 598 PGGPPR--PPGPPP-PP-------GTPRAPGA---PPPPPPP---GGLKKMQTT--KKGGVQRAPDVIQLYQKLFKRDGS
KfCHUP1 754 KAKLVG--PEGAPP-PP-------PPPPGKGG--PPPPPPPP---GAPGGKLAPIHTKDGVQRAPEVIQLYQSLM-RIGR
OsCHUP2 504 ---------SAANP-PK-----------------PPPPPPPP---KFSTRN------AGVMKRAPQVAELYHSLMRRDSK
NnCHUP2 497 --------PAPPPP-PP-----------------PPPPPPPP---KFSIGS-----STSLVQRAPQVVEFYHSLMKRESR
AtrCHUP2 564 ---------KVAPP-PP-----------------PPPPPPPP---KFSMRS-----SASVMQRAPEVVEFYHSLMKRDGR
PpCHUP1c 931 VAGPLG---PPPPP-PP-------PPPLTPGALCPPPPPPTP---GSLIKG-SG---AEKMQRAPGVVEFYQSLMKRDAK
PpCHUP1a 805 GGGV----PPPPPP-PP-------PPRGGPGA--PPPPPPPPPM-GGLSKM-GK--KTDDVHRAPEVVEFYQSLMKRDAK
PpCHUP1b 831 PGGFVI---PPPPP-R------------GPGA--LLPPPPPPSLKGSLSRTQGN--HSDDVHRAPEVVEFYHSLMKRDSK
MpCHUP1 743 PP------PPPPPP-PP-------RLPGAPGAPPPPPPPPPP---GGKLGA-SV--GGNKMQRAPEVVEFYQSLMKIGAK
OsCHUP1 615 PPPPR---PPGAPP-PP-------PPPGKPGG--PPPPPPRP---GSLPRN-LA--GGDKVHRAPEVVEFYQSLMKREAK
AtCHUP1 669 SARPPL--PGGGPP-PPPPPPGGGPPPPPGGG--PPPPPPPP---GALGRG-AG--GGNKVHRAPELVEFYQSLMKRESK
AtrCHUP1 698 VAPPPL--PPGAPP-PP-------PPPPLSGA--PPRPPPPP---GSLPKG-SG---GDKVHRAPELVEFYQSLMKREAK
NnCHUP1a 686 A------------P-PP-------LSPP--GE--PPCPPPPP---GSLPGG-SS--TGDKVHRAPEIVEFYRTLMKREAK
NnCHUP1b 684 APPP----PPGAPP-PP-------PPPP--GG--PPRPPPPP---GSLPRG-SG--TGDKVHRAPELVEFYQTLMKREAK
SmCHUP1a 609 PPRLAGTAPPPPPP-PPGIPGAPPLPPGAPGA--PPPPPPLP---GMGKPQ-GQ--SGSKVQRAPEVVEFYQSLMKRDAR
SmCHUP1b 612 PPRLAGTAPPPPPP-PPGIPGAPPLPPGAPGA--PPPPPPLP---GMGKPQ-GQ--SGSKVQRAPEVVEFYQSLMKRDAR
AcCHUP2 720 SAGPPL--PPPPPP-PP--------PPRAPGA---PPPPPLP---GSLKVQ-GT--GKEQVQRAPEVVEFYQSLMRREAK
AcCHUP1 737 G--------APPPP-PP--------PPRAPGA--PPPPPPPP---GSLKSQ-GA--SGDKVQRAPEVVEFYQSLMRREAK
LjCHUP1 721 ARGPASALPGPPPPPPP--------PPRVPGA--PPPPPPPP---GSLKSQ-GP--TGDKVQRAPEVVEFYQSLMRREAK
MvCHUP1 660 AKLA---VKKDDDGGHDARSEMVNEIEGRSSHLMEVKQDVEVQGEFINYLATEVTGLKIRDMDKVVLFVEWLDNELSYLV
KfCHUP1 818 GYTGEGGISGPGMDVDGARGEMVAEIENRSSHLLAIKADVEAQADFVKSLAQEVRAAEYDRIEDVVAFVSWLDEELSFLV
OsCHUP2 548 KDTSGSGI-CETANSANVRSSMIGEIENRSSHLQAIKADVETQGEFVKSLIKEVTNAAYKDIEDVVAFVKWLDDELGFLV
NnCHUP2 543 RESSSGGI-CDAPDVANVRSSMIGEIENRSSHLLAIKADVETQGEFVNSLIREVNNAVYQDIEDVVAFVKWLDDELCFLV
AtrCHUP2 609 KDSAAGGV-CDAPGVTNVHSSMIGEIENRSSYLLAIKADVETQGEFVRSLIREVNNAVYRDIEDVVAFVKWLDDELCFLV
PpCHUP1c 993 QSLS---SPGGTVSNSEARNNIIGEIENRSTHLLAIKADVETQGEFVESLAAEVRAASFSNIEEVVEFVVWLDEELSFLV
PpCHUP1a 867 SAV----VNTAGGNNPEARNNMIGEIENRSTHLLAIKADVETQGEFVMSLAAEVRAAVYGDIKDVVEFVNWLDEELSFLV
PpCHUP1b 891 SAVSN----SGGGTDPTARNNMIGEIENRSAHLLAIKADVETQGDFVMSLAVEVRAAEFTDIEDVVNFVRWLDDELSYLV
MpCHUP1 803 KE-----LAASGGNSAEARNNMVGELENRSAHLLAIKADVETQADFVKSLATEVRGAQYTDIEDVVAFVTWLDDELSFLV
OsCHUP1 676 KDTTS--LGSTTSSVSDVRSNMIGEIENRSTFLLAVKVDVETQGDFVESLANEVRAASFVNIDDVVAFVNWLDEELSFLV
AtCHUP1 738 KEGAPSLISSGTGNSSAARNNMIGEIENRSTFLLAVKADVETQGDFVQSLATEVRASSFTDIEDLLAFVSWLDEELSFLV
AtrCHUP1 759 KDGSS--VASSTSNTADVRSNMIGEIENRSTFLLAVKADVETQGDFVQSLATEVRAATFNNIEDVLSFVNWLDEELSFLV
NnCHUP1a 736 KDTSS--LTLFTSDASDTRSNMIGEIENRSAFLLSVRADVETQGDFVQSLATEVQAASFTNIEDLMLFVNWLDEELSFLV
NnCHUP1b 742 KDTST--LTSFTPNTSDVRSNMIGEIENRSSFLLAVKADVETQGDFVQSLATEVRAASFTNIEDLVSFVNWLDEELSFLV
SmCHUP1a 680 KDAA---VSSSGNASSEARSNLIGEIENRSSHLLAIKADVETQGDFVNSLAAEVRAAVYSNIDDILAFVNWLDEELAFLV
SmCHUP1b 683 KDAA---VSSSGNASSEARSNLIGEIENRSSHLLAIKADVETQGDFVNSLAAEVRAAVYSNIDDILAFVNWLDEELAFLV
AcCHUP2 780 KDTS---LGASDVNASDARNNMIGEIENRSAFLLAVKADVETQGDFVQSLATEVREAAYTDIEDVIAFVAWLDEELSFLV
AcCHUP1 792 NNTS---VGATDVNVSDARNNLIGEIENRSAFLLAVKADVETQGEFVQSLAAEVRDAAYTDIEDVVAFVSWLDEELSFLV
LjCHUP1 785 KDTG---LGAADANVSDVRNNMIGEIENRSAFLLAVKADVETQGDFVNSLAAEVRNAAYTDIEDVVAFVTWLDEELSFLV

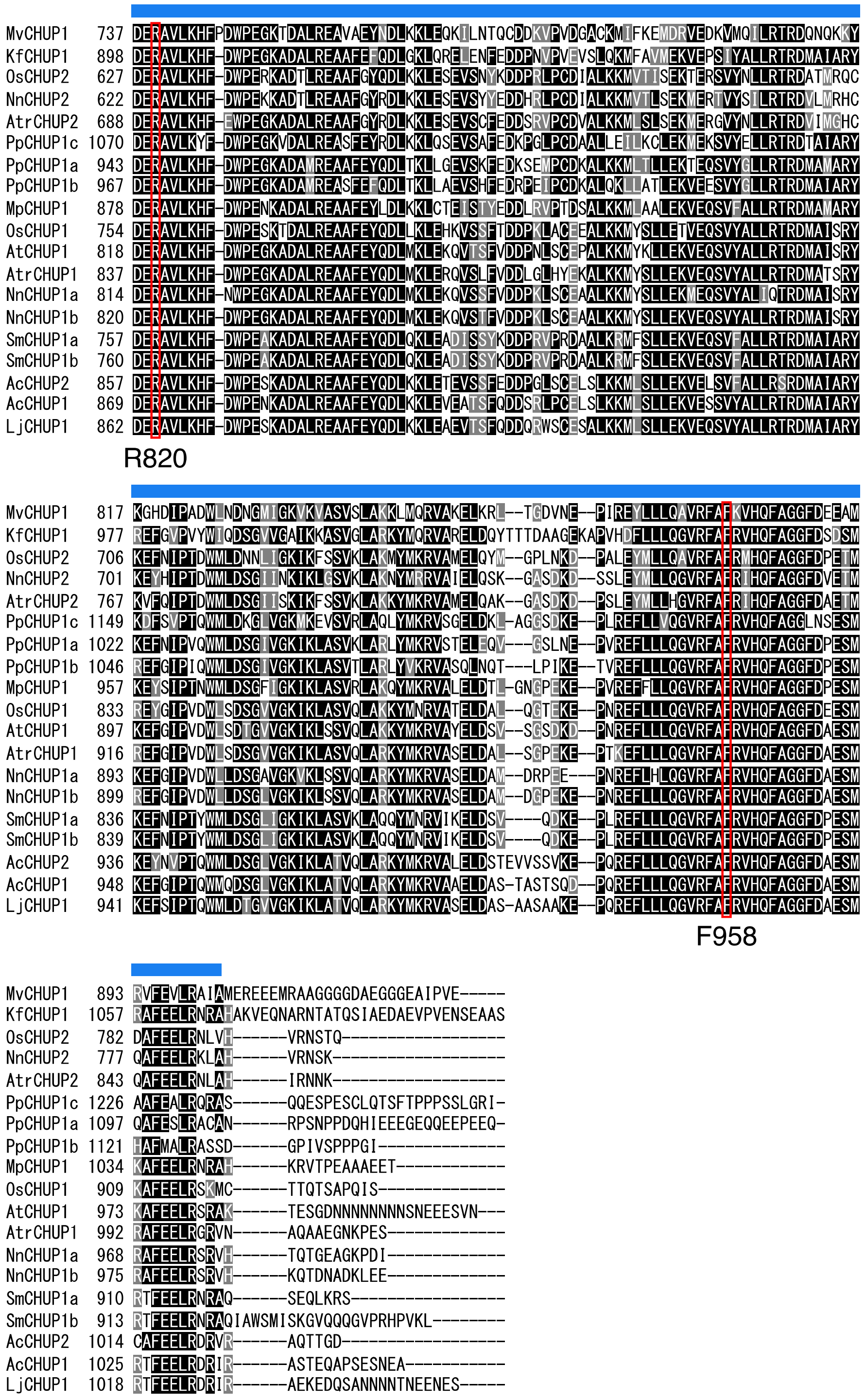


**Figure S6. Protein sequence alignment of the C-terminal region of CHUP1**

All aligned sequences were obtained from GenBank. Green bar, profilin-rich region of the FH1-like domain. Blue bar, FH2-like domains conserved in CHUP1 family proteins. Red boxes, R820 and F958.

Ac, *Adiantum capillus-veneris* (Ferns); At, *Arabidopsis thaliana* (Angiosperms); Atr, *Amborella trichopoda* (Angiosperms); Kf, *Klebsormidium flaccidum* (Klebsormidiales); Lj, *Lygodium japonicum* (Ferns); Mp, *Marchantia polymorpha* (Liverworts); Mv, *Mesostigma viride* (Mesostigmatales); Nn, *Nelumbo nucifera* (Angiosperms); Os, *Oryza sativa* (Angiosperms); Pp, *Physcomitrella patens* (Mosses); Sm, *Selaginella moellendorffii* (Lycophytes).

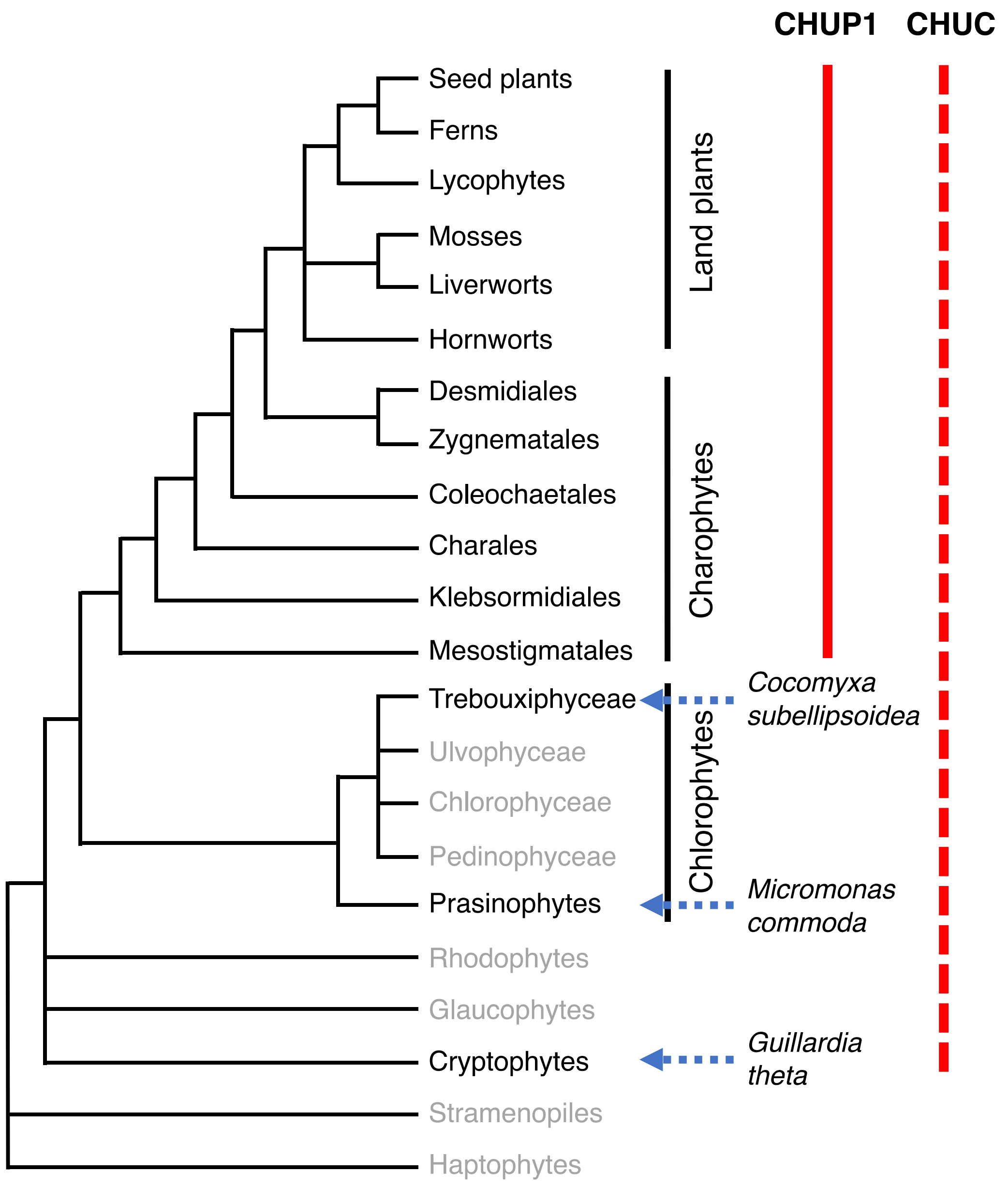


**Figure S7. Organismal lineages and distributions of CHUP1 and CHUC genes**

The topology of lineages is adapted from (Li et al., 2015). CHUP1 genes were identified in the genome and/or transcriptome databases of charophytes and land plants but not in those of chlorophytes. By contrast, CHUC genes were identified in unicellular algae such as *Guillardia theta*, *Micromonas commoda*, and *Coccomyxa subellipsoidea* as well as in land plants.

### Supplemental Tables

**Table S1.** **Data collection and refinement statistics**

| Statistics | CHUP1_756-982/ C864S (SeMet) | CHUP1_716-982 (Native) |
|---|---|---|
| **Data collection** | | |
| Space group | $P6_222$ | $I2_13$ |
| Cell dimensions | | |
| *a*, *b*, *c* (Å) | 123.3, 123.3, 160.8 | 180.7, 180.7, 180.7 |
| α, β, γ (°) | 90.0, 90.0, 120.0 | 90.0, 90.0, 90.0 |
| Resolution (Å) | 50.0–2.8 (2.85–2.80) | 50.0–3.0 (3.05–3.00) |
| $R_{merge}$ | 0.201 (>1) | 0.114 (>1) |
| $I / \sigma I$ | 27.2 (2.2) | 30.6 (2.2) |
| Completeness (%) | 99.9 (100.0) | 99.9 (100.0) |
| Redundancy | 41.9 (43.8) | 21.7 (22.8) |
| **Refinement** | | |
| Resolution (Å) | 50.0–2.8 | 50.0–3.0 |
| No. reflections | 18,407 | 19,727 |
| $R_{work}$ / $R_{free}$ | 0.202 / 0.228 | 0.174 / 0.224 |
| No. atoms | | |
| Protein | 1,815 | 3,791 |
| Water | 7 | 0 |
| B-factors (Å$^2$) | | |
| Protein | 72.0 | 94.8 |
| Water | 57.5 | |
| Rms deviations | | |
| Bond lengths (Å) | 0.009 | 0.010 |
| Bond angles (°) | 1.09 | 1.29 |

Values in parentheses are for highest-resolution shell.

The $R_{free}$ value was calculated for the $R$ factor using a test set (5%) of reflections not used in the refinement.

**Table S2. Binding of CHUP1-C to skeletal muscle actin filaments**

| | control | + shearing | + gelsolin | + EGTA |
|---|---|---|---|---|
| Fraction of CHUP1-C in ppt (%) Average ± SD | 4.7±1.4 | 10.4±2.9 | 5.1±1.8 | 5.7±5.4 |
| N | 6 | 6 | 3 | 3 |

Data shown in Figure S2A were analyzed using ImageJ (https://imagej.nih.gov/ij/), and the fraction of CHUP1-C in the pellet after ultracentrifugation was calculated from Sample 1 (control), Sample 2 (+ shearing), Sample 3 (+ gelsolin), and Sample 4 (+ EGTA). The difference between control and + shearing was statistically significant according to a Student's *t*-test ($p$=0.007).
